# A non-canonical sensing pathway mediates *Plasmodium* adaptation to AA deficiency

**DOI:** 10.1101/2022.12.01.518651

**Authors:** Inês M. Marreiros, Sofia Marques, Ana Parreira, Vincent Mastrodomenico, Bryan C. Mounce, Chantal T. Harris, Björn F. Kafsack, Oliver Billker, Vanessa Zuzarte-Luís, Maria M. Mota

## Abstract

Eukaryotes have canonical pathways for responding to amino acid (AA) availability. Under AA-limiting conditions, the TOR complex is repressed, whereas the sensor kinase GCN2 is activated. While these pathways have been highly conserved throughout evolution, malaria parasites are a rare exception. Despite auxotrophic for most AA, *Plasmodium* does not have either a TOR complex nor the GCN2-downstream transcription factors. While Ile starvation has been shown to trigger eIF2α phosphorylation and a hibernatory-like response, the overall mechanisms mediating detection and response to AA fluctuation in the absence of such pathways has remained elusive. Here we show that *Plasmodium* parasites rely on an efficient sensing pathway to respond to AA fluctuations. A phenotypic screen of kinase knockout mutant parasites identified nek4, eIK1 and eIK2 – the last two clustering with the eukaryotic eIF2α kinases - as critical for *Plasmodium* to sense and respond to distinct AA-limiting conditions. Such AA-sensing pathway is temporally regulated by these kinases at distinct life cycle stages and allows parasites to actively fine-tune replication and development in response to AA availability. Collectively, our data identify a previously unknown set of heterogeneous responses to AA depletion, mediated by a complex mechanism that is critical for modulating parasite cell cycle and survival.

---

Malaria, a disease caused by *Plasmodium* parasites, kills a child every minute. While there have been significant reductions in the number of infection and mortality rates since 2000 (51% and 27% reduction, respectively), the plateau in progress, first observed in 2017, still persists. Notably, during the past two years, the COVID-19 pandemic has resulted in unprecedent challenges to health systems worldwide, having disrupted malaria services, leading to a marked increase in cases and deaths. In 2020, there were an estimated 241 million malaria cases and 627 000 malaria deaths worldwide, which represents about 14 million more cases and 69 000 more deaths compared to 2019^1^.

Both parasite and host genetics, as well as host immune responses, contribute to the outcome of infection^2–5^. Still, the most consistent indicator of disease severity is parasite biomass^6^. *Plasmodium* is a rapidly multiplying unicellular organism undergoing a complex developmental cycle that takes place in the mammalian and mosquito hosts – a lifestyle that requires rapid adaptation to distinct environments. In the mammalian host, *Plasmodium* initially infects hepatocytes to generate thousands of merozoites, which are later released into the bloodstream where they invade and replicate within red blood cells (RBCs). Each merozoite replicates by schizogony to generate 10-30 new merozoites every cycle, which occurs at 24, 48 or 72 hours - depending on the *Plasmodium* species. As a rapidly multiplying obligatory intracellular parasite, *Plasmodium* has high nutritional demands relying on the nutrients provided by its host to survive. To cope with the different environments and conditions its life cycle entails, *Plasmodium* mainly relies on the expression of distinct transcriptomes at each developmental stage^7–9^. However, besides the ever-changing environment accompanying its normal life cycle progression, the environment within a single host is also extremely dynamic, with parasites constantly facing variable immune responses, drug pressures and metabolic fluctuations^10^. We and others have provided evidence that parasite adaptation to unpredictable fluctuations within each niche requires alternative strategies, such as efficient sensors, and a tight regulation of nutrient-sensing signaling pathways^11–13^. *Plasmodium* parasites are auxotrophic for most amino acids (AA), and the blood stages acquire the majority of them through the digestion of host erythrocyte hemoglobin^14,15^. However, some AA are either absent (isoleucine) or rare (methionine) within human hemoglobin and must be acquired from extracellular sources^16,17^.

In eukaryotes, from yeast to mammals as well as plants, the target of the rapamycin (TOR) complex and the general control nonderepressible 2 (GCN2)/eIF2α signaling cascades are two well-characterized mechanisms to sense AA fluctuations^18,19^. Notably, no homolog of the TOR complex has been identified to date in *Plasmodium;* however, a downstream effector, *Maf1*, is conserved in *P. falciparum* and has been reported to be essential for the asexual blood stage of the parasite^20^. On the other hand, although the *Plasmodium* eIK1 kinase^21^ closely clusters with the GCN2 AA sensing-kinase, the parasite lacks some key downstream components of this pathway, including the downstream transcription factors as well as the biosynthetic pathways that mediate GCN2 action - namely the orthologues for starvation-response regulators such as GCN4 in yeast and ATF4 in mammals^22,23^.

This has led some to propose that *Plasmodium* evolved a stripped-down starvation-response pathway^24^, implying a reduced capacity to adapt to AA depletion, rather than actively responding to alterations in AA levels. The present paradigm is that *P. falciparum*, in the absence of canonical eukaryotic nutrient stress-response pathways, can cope with an inconsistent AA supply by hibernating until more nutrients are provided^24^. We now challenge this paradigm by showing that both *P. berghei* and *P. falciparum* parasites can actively mount a response to a decrease in two distinct AAs – methionine and isoleucine – by activating in each life cycle stage a different protein kinase, ultimately leading to a reprograming of replication and parasite development.

## AA depletion impacts *Plasmodium* cell cycle pace and replication rates

To assess the effect of AA depletion during parasite intra-erythrocytic development, we monitored the growth of both *P. falciparum* and *P. berghei* parasites cultured in media with or without methionine (Met) or isoleucine (Ile). Sorbitol-synchronized *P. falciparum* NF54 wild-type (WT) parasites cultured in medium containing methionine (MetS) or isoleucine (IleS) (0.100 mM MetS; 0.38 mM IleS) develop and replicate every 48h, resulting in a marked increase in parasitemia with each generation (**Figure 1a**). However, when cultured in either Met deficient (MetD) or Ile deficient (lleD) medium, parasites undergo impaired growth, as evidenced by a marked reduction in parasitemia **(Figure 1a**). Notably, while the effect of IleD on parasitemia is evident from day 2 onwards, parasitemia in MetD only becomes significantly different from day 4 onwards **(Figure 1a)**. Such difference may be explained by the fact that Ile is totally absent from human hemoglobin, while Met is present, although at low levels^25^. Neither IleD^24^ nor MetD impact parasite viability for a 48h period, as synchronized ring-stage forms quickly resume growth upon AA supplementation **(Figure 1 - Figure supplement 1a).** Methionine, among the twenty amino acids, plays an integral role in protein biosynthesis and in regulation of translation and global gene expression - as it is coded by the translation initiation codon. Met is metabolized into S-adenosylmethionine (SAM), the cellular methylation currency, via an ATP-dependent process which is catalyzed by the *Plasmodium* SAMS enzyme. Thus, besides MetD we have also focused on the SAMS enzyme, as SAM is the major biological methyl donor and also the precursor of polyamines, via the aminopropylation pathway, or glutathione (GSH), when entering the transsulfuration pathway, whereby homocysteine (Hcy) is hydrolyzed into GSH (**Figure 2a)**. Addressing both MetD conditions and SAMS cKD parasites allowed us to establish the role of Met in protein incorporation but also to disentangle the role of methylation, GSH and polyamines in parasite growth.

**Figure 1.**
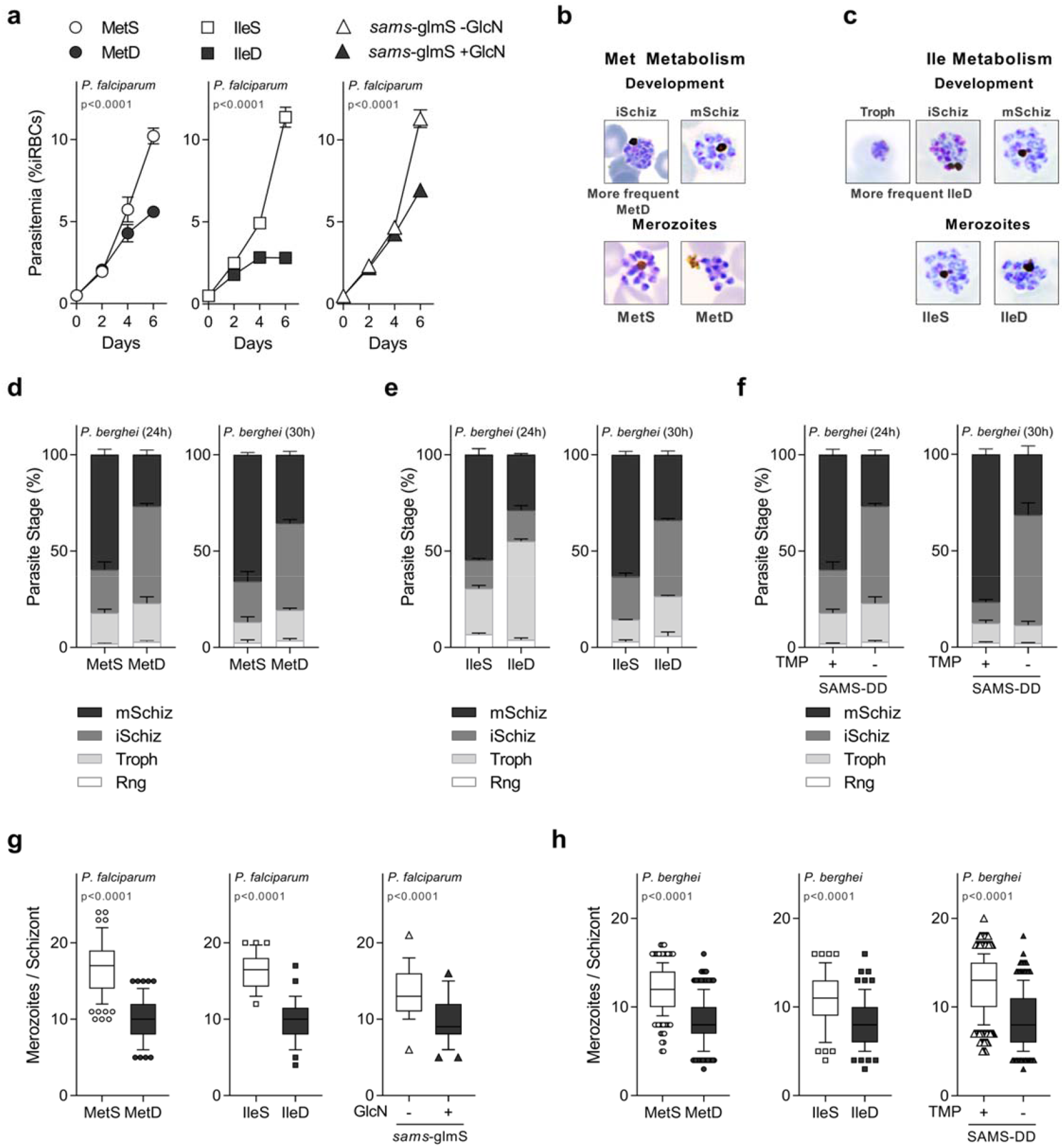
AA availability impacts parasite intra-erythrocytic development and replication. **(a)** Parasitemia of synchronized *P. falciparum* NF54 wt parasites cultured in AA deficient media (MetD, IleD), or *P. falciparum* parasites knock-down for the SAMS enzyme (*sams*-glmS + GlcN). **(b, c)** Representation of blood-stage development and replication in *P. berghei* parasites upon AA withdrawal from the culture media. Parasites were cultured in **(b)** MetD- or **(c)** IleD-media for 30h and visualized by microscopy after Giemsa staining. **(d-f)** Quantification of *P. berghei* intra-erythrocytic developmental stages, by microscopy analysis after *in vitro* culturing for 24h or 30h in **(d)** MetD- or **(e)** IleD-media and in **(f)** parasites knock-down for the SAMS enzyme (SAMS-DD - TMP). **(g, h)** Box plot of mean merozoites number per schizont in **(g)** *P. falciparum* or **(h)** *P. berghei* parasites, cultured in MetD-, IIeD-media or in SAMS cKD parasites. **a.** Data represents the mean percentage of iRBCs ± SEM (2-way ANOVA), determined in 2 independent experiments, each performed at least in triplicate. **d-f.** Parasite staging was performed in triplicate, averaged and repeated at least in 2-3 independent experiments. Data is shown as mean percentage of parasites at each developmental stage with error bars representing SEM. **g, h.** Data is represented as box-whisker plot of mean merozoite number per schizont ± SD (Mann-Whitney), with the median represented at the center line. Boxplots show the data of 3-5 independent experiments for the following number of schizonts, *P. falciparum*: MetS, *n*=107, MetD, *n*=86; IleS, *n*=32, IleD, *n*=37; *sams*-glmS -GlcN, *n*=25; *sams*-glmS +GlcN, *n*=45; *P. berghei*: MetS, *n*=235; MetD, *n*=275; IleS, *n*=65; IleD, *n*=70; SAMS-DD + TMP, *n*=405, SAMS-DD - TMP, *n*=325. **Figure 1 – Source data 1** *P. falciparum* attenuated growth under low AA levels or in SAMS knock-down parasites related to Figure 1a. **Figure 1 – Source data 2** Cell cycle progression and *in vitro* maturation rates of *P. berghei* parasites growing under AA deficiency or lacking the SAMS enzyme related to Figure 1d-f. **Figure 1 – Source data 3** Impact of AA depletion in *P. falciparum* and *P. berghei* replication rates related to Figure 1g,h.

**Figure 2.**
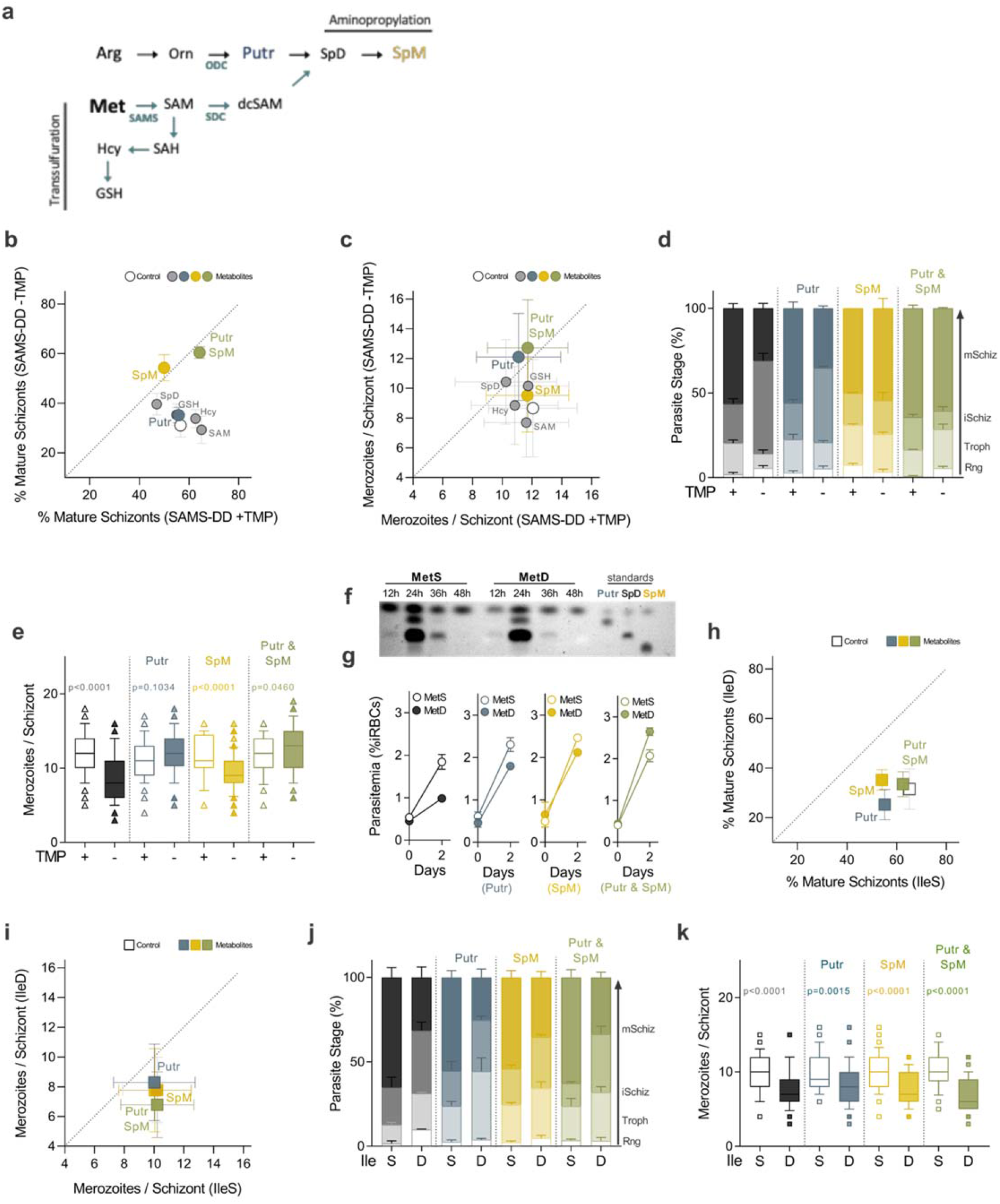
Polyamines regulate parasite’s response to methionine depletion. **(a)** Schematic representation of the close connection between the methionine cycle and the aminopropylation pathway in malaria parasites, with both pathways relying on the *Plasmodium* bifunctional SDC/ODC enzyme. **(b)** Percentage of mature segmented schizonts and **(c)** merozoite number per schizont in *Pb*SAMS-DD -TMP parasites cultured for 24h in media supplemented with each SAM-dependent metabolite. **(d)** Quantification of *P. berghei* intra-erythrocytic developmental stages in *P. berghei* parasites knock-down for the SAMS enzyme, *in vitro* cultured for 24h in media supplemented with the polyamines Putr, SpM or both. **(e)** Box plot of mean merozoite numbers of *Pb*SAMS-DD parasites cultured as in d. **(f)** Thin-layer chromatography (TLC) analysis on synchronized *P. falciparum* NF54 WT parasites cultured in MetS or MetD-media. Polyamine levels were assessed at 12h (ring-stage), 24h (trophozoite-stage), 36h (schizont stage) and 48h (next generation ring-stages) of *in vitro* culture. The same number of parasites was loaded per condition and timepoint. Due to its very low abundance in the parasite, SpM could not be detected by TLC analysis. **(g)** Parasitemia in synchronized *P. falciparum* 3D7 WT parasites cultured in MetD-media supplemented with the polyamines Putr, SpM or both **(h)** Percentage of mature segmented schizonts and **(i)** mean merozoite numbers in *P. berghei* WT parasites cultured for 24h in IleD- or IleD-media supplemented with each polyamine. **(j,k)** Quantification of **(j)** *P. berghei* intra-erythrocytic developmental stages and **(k)** box plot of mean merozoite numbers of *P. berghei* WT parasites, *in vitro* cultured for 24h in IleD- or IleD-media supplemented with polyamines. **b, c, h, i.** All media supplementations were performed in triplicate and averaged, with each data point representing the **(b, h)** mean percentage of mature schizonts or **(c, i)** the mean merozoite number produced per schizont and error bars representing the SE Diff or SD between replicates, respectively. Representative results of 2-3 independent experiments are shown. **d, j.** Parasite staging was performed in triplicate, averaged and repeated in 2-3 independent experiments. Data is shown as mean percentage of parasites at each developmental stage, with error bars representing SEM. **e, k.** Data is represented as box-whisker plot of mean merozoite number per schizont ± SD (2-way ANOVA), with the median represented at the center line. Boxplots show the data of 2 independent experiments and for the following number of schizonts, SAMS-DD +TMP, *n*= 122, SAMS-DD -TMP, *n*= 117; SAMS-DD +TMP +Putr, *n*= 50, SAMS-DD -TMP +Putr, *n*= 48; SAMS-DD +TMP +SpM, *n*= 37, SAMS-DD -TMP +SpM, *n*= 102; SAMS-DD +TMP +Putr & SpM, *n*= 47, SAMS-DD -TMP +Putr & SpM, *n*= 47;: IleS, *n*= 38, IleD, *n*= 47; IleS +Putr, *n*= 59, IleD +Putr, *n*= 66; IleS +SpM, *n*= 56, IleD + SpM, *n*= 57; IleS +Putr & SpM, *n*= 38, IleD +Putr & SpM, *n*= 53. **g.** Data represents the mean percentage of iRBCs ± SEM (2-way ANOVA), determined in 2 independent experiments, each performed at least in triplicate. **Figure 2 – Source data 1** Screen of the methionine-downstream metabolites in schizont maturation and replication rates of parasites lacking the SAMS enzyme related to Figure 2b,c. **Figure 2 – Source data 2** *In vitro* maturation and replication rates of *P. berghei* parasites lacking the SAMS enzyme upon polyamines supplementation related to Figure 2d,e. **Figure 2 – Source data 3** *P. falciparum* growth in MetD media supplemented with polyamines related to Figure 2g. **Figure 2 – Source data 4** Screen of the methionine-downstream metabolites in schizont maturation and replication rates of parasites growing under IleD conditions related to Figure 2h,i. **Figure 2 – Source data 5** *In vitro* maturation and replication rates of *P. berghei* parasites growing in IleD media supplemented with polyamines related to Figure 2j,k.

To do so, we employed a previously generated^26^ glucosamine (GlcN)-regulatable parasite line in which the *P. falciparum* SAMS, the first and rate-limiting enzyme of the methionine cycle, was conditionally knocked down (*Pfsams-glmS* parasite line). Interestingly, the knockdown of the SAMS enzyme, achieved by GlcN addition to the culture media (*Pfsams*-glmS +GlcN), phenocopies WT parasites growing in MetD media, as evidenced by a marked reduction in parasitemia from day 4 onwards **(Figure 1a)**.

We next sought to determine whether reduced parasitemia observed upon AA limitation was an outcome of defective parasite development. We employed the *P. falciparum* 3D7 stain which exhibited a shorter life cycle length, replicating every 36h to 42h in control conditions - in accordance to previous studies estimating a 38.8h cell cycle length for this particular strain^27,28^. Time course analysis of synchronized *P. falciparum* 3D7 ring-stages growing in MetD or IleD-media reveals impaired parasite development in both conditions, yet with some particularities unique to each condition **(Figure 1 - Figure supplement 1b,c).** Both MetD- and IleD-media growing parasites fail to complete maturation in a timely and coordinated manner, arresting at the immature schizont-stage, as evidenced by an increase in the time spent at this stage **(Figure 1 - Figure supplement 1b,c).** However, Ile depletion has an even more pronounced impact in cell cycle progression, evidenced by a reduction in cell cycle pace earlier in development, at the trophozoite stage, with parasites arresting for one complete developmental cycle at this particular stage **(Figure 1 - Figure supplement 1c).** Indeed, such a striking delay in progression of isoleucine-starved parasites through the trophozoite stage was first documented by Babbit *et al*. which proposed a dormancy-like model upon AA starvation^24^. This prompt and sharp impact of Ile depletion on parasite growth – in comparison to methionine depletion – might result from a quicker depletion of Ile stocks, as it is totally absent from human Hb. However, our results show that even though the appearance of next generation merozoites is delayed both in MetD and IleD conditions, parasites can invade erythrocytes efficiently, as evidenced by the emergence of second-generation ring-stage forms that peak at ~72h-84h of development **(Figure 1 - Figure supplement 1b,c).**

Notably, when employing a malaria parasite of rodents, *P. berghei* ANKA, a similar impact was observed in MetD and IleD conditions **(Figure 1b, c)**. Blood stages of rodent malaria parasites can only be maintained in culture for one developmental cycle as newly developed merozoites cannot reinvade erythrocytes anew. Even though synchronization cannot be achieved in this short-term *ex vivo* systems, parasites fully mature from rings/young trophozoites into segmented schizonts yet, even when fully mature, schizonts do not rupture *in vitro*^29^ - with cultures being enriched for this particular stage, allowing a thorough characterization of schizont development.

Microscopy analysis show that *P. berghei* parasites either growing in MetD or IleD media exhibit a delay in maturation *in vitro*, with only 30% of the parasites progressing through the cycle at a normal pace and successfully developing into mature schizonts **(Figure 1d, e).** As for *P. falciparum*, while IleD leads to a significant delay in both trophozoite- and immature schizont stages (**Figure 1e)**, MetD mainly impacts the immature-to-mature schizont transition **(Figure 1d).** Surprisingly, providing *P. berghei* parasites an additional 6h of maturation time (30h of *in vitro* culture) did not reflect in increased maturation rates under AA deficiency. This suggests that ~70% of parasites growing under AA-limiting conditions exhibit a growth arrest which is reminiscent of the hibernatory phenotype previously described by *Babbit* et al^24^. Again, a *P. berghei* transgenic parasite line (*Pb*SAMS-DD parasite line) in which the first enzyme of the methionine cycle was knocked down (*Pb*SAMS-DD -TMP) phenocopied WT parasites cultured in MetD conditions (**Figure 1f).** In this species, SAMS knockdown was achieved by employing the destabilizing domain (DD) system^30^. The construct containing the *sams* gene fused to the DD and the hemagglutinin (HA) tag was introduced into *P. berghei* genome using double crossover homologous recombination (**(Figure 1 - Figure supplement 1d,e).** Protein stabilization *in vivo* was achieved by providing trimethoprim (TMP) in drinking water for 2 days prior to infection (*Pb*SAMS-DD +TMP), while SAMS conditional knockdown (*Pb*SAMS-DD -TMP) was achieved in non-TMP treated mice, as evidenced by a significant decrease in SAMS-HA-DD protein levels, measured by immunoblotting analysis **(Figure 1 - Figure supplement 1f).** Similar results were observed by immunofluorescence analysis of *P. berghei* trophozoites and schizonts in which the SAMS enzyme was knocked down (*Pb*SAMS-DD -TMP; Troph, Schiz), as evidenced by a significant decrease in SAMS-HA-DD protein levels relative to control parasites **(Figure 1 - Figure supplement 1g).** Notably, *Pb*SAMS-DD -TMP merozoites exhibit SAMS expression levels per merozoite comparable to those observed in control parasites **(Figure 1 - Figure supplement 1h).** *Plasmodium* schizogony culminates in the production of individualized merozoites that quickly re-invade new erythrocytes. We observed that under AA deficient conditions, ~30% of parasites successfully complete the cell cycle in a timely manner, developing into mature schizonts containing individualized next generation merozoites. Yet, we observed that the number of merozoites each schizont contained was dependent on the growth conditions. Microscopy and flow cytometry analysis in *P. falciparum* and *P. berghei* showed significantly fewer merozoites in schizonts grown in MetD or IleD media, as well as in transgenic parasites knockdown for the SAMS enzyme (*Pfsams-glmS* +GlcN*; Pb*SAMS-DD -TMP) **(Figure 1g, h; Figure 1 - Figure supplement 2a-d).** Again, allowing *P. berghei* parasites an additional 6 h of maturation time (30h of *in vitro* culture) did not restore merozoite numbers in any of the conditions **(Figure 1 - Figure supplement 2e),** suggesting that lower replication levels result neither from the asynchronous nature of *Plasmodium* schizogony nor from a delayed maturation, but rather from an ability of parasites to adapt to low AA levels. To disclose the role of AA metabolism in the dynamics of an *in vivo* malaria infection, we provided a short-term, methionine-deprived (0% methionine, MetD) regimen to BALB/c mice for 3 weeks prior to a *P. berghei* ANKA infection. Body weight analysis shows that, in accordance to previous findings^31^, a MetD regimen leads to a 30% weight loss relative to the methionine-sufficient (MetS) regimen **(Figure 1 - Figure supplement 2f).** Flow cytometry analysis of MetD-fed BALB/c mice infected with WT parasites shows a significant reduction in peripheral parasitemia relative to the MetS-fed group **(Figure 1 - Figure supplement 2g)**. Interestingly, the same was observed in mice fed on a MetS diet but infected with *Pb*SAMS-DD -TMP parasites. **(Figure 1 - Figure supplement 2h).** Moreover, the reduced number of merozoites per schizont we observed after conditional knockdown of *Pb*SAMS *in vitro* was also confirmed *in vivo* **(Figure 1 - Figure supplement 2i).** Altogether, these findings show that fluctuations in AA availability affect parasite growth, by either stalling progression through the intra-erythrocytic asexual cell cycle or by reducing parasite replication rates.

## Polyamines regulate *Plasmodium* response to MetD

We next sought to identify which signal functions as environmental cue for a parasite stress response to AA deficiency. While Ile shares the first enzymatic steps of its metabolism with two other AA (leucine and valine)^32^, the initial steps of Met metabolism are unique among all AAs, rendering the products of its metabolism ideal candidates. To find out which Met-downstream metabolites could rescue attenuated replication and delayed maturation in *P. berghei* parasites knockdown for the SAMS enzyme, we supplemented media with SAM, Hcy, GSH or the polyamines putrescine (Putr), spermidine (SpD) and spermine (SpM) (**Figure 2a).**

Microscopic analysis shows that SAM rescues neither impaired maturation, nor reduced merozoite numbers produced by *Pb*SAMS-DD -TMP schizonts, relative to the control group **(Figure 2 - Figure supplement 1).** This is not surprising, since SAM transport across the plasma membrane occurs at a minimal extent in mammalian cells^33^. However, the results show that among the SAM-downstream metabolites, polyamines, namely Putr and SpM, were the sole metabolites regulating parasite response to SAMS deficiency **(Figure 2b-e, (Figure 2 - Figure supplement 1a,b**). In *Pb*SAMS-DD schizonts knockdown for the SAMS enzyme (-TMP), while SpM reverses the impaired maturation, Putr fully restores merozoite numbers **(Figure 2b-e).** Additionally, concomitant addition of SpM and Putr rescues not only the delayed development of *Pb*SAMS-DD -TMP parasites but also the reduced replication rates – as evidenced by a higher percentage of mature schizonts and by an increased merozoite number **(Figure 2b-e).** This data further corroborates the existence of two distinct parasite populations – one that stalls development and other that, albeit completing the cycle at a normal pace, develops into schizonts containing fewer progeny – which are independently regulated by distinct metabolites (SpM or Putr) and likely, by distinct mechanisms.

In an attempt to unveil how parasite replication and development are independently regulated by distinct polyamines, we assessed the impact of MetD on polyamine biosynthesis throughout parasite intra-erythrocytic development. Thin layer chromatography (TLC) analysis of *P. falciparum* NF54 parasites shows an overall reduction in polyamine synthesis throughout development in MetD **(Figure 2f).** Notably, the effect of MetD on Putr production is more striking during early stages of parasite development (trophozoite, 24h) whereas the production of SpD - and likely that of its downstream product, SpM - is mainly reduced during late schizogony (36h). This supports our previous findings in the rodent malaria species, *P. berghei* ANKA, showing a role for Putr during replication **(Figure 2e)**, and for SpM at later stages of parasite development, particularly during schizont segmentation **(Figure 2d)**. These findings were recapitulated in the human malaria parasite, *P. falciparum*, in MetD conditions, as evidenced by a sharp increase in parasitemia upon polyamine supplementation **(Figure 2g).** Indeed, our data is in agreement with previous findings showing that inhibition of *P. falciparum* ODC enzyme (critical for Putr synthesis) induces a cytostatic arrest at the trophozoite stage, halting parasite development at G1/S transition – a phenotype that is rescued by the addition of exogenous polyamines^34–36^. So far, our data show that the developmental and replication responses to methionine depletion are independent, each relying on a different polyamine. Not surprisingly, the effects of Met-downstream metabolites did not extend to conditions of Ile depletion, showing that these were specific to Met depletion signaling **(Figure 2h-k).** The data show that separate mechanisms are orchestrated upon depletion of each amino acid (Met or Ile), with the downstream metabolites triggering these responses being specific to the metabolic pathway that is being affected. It is important to note that the differential responses observed at the phenotypic level likely translate the distinct roles each AA plays in the cell. Our findings so far led us to hypothesize that *Plasmodium* parasites have the ability to sense fluctuations in AA levels - as well as in the downstream metabolites – which independently regulate parasite development or replication, functioning as key molecular players during distinct stages of parasite’s life cycle.

## Nek4, eIK1 and eIK2 are key regulators of *parasite* response to AA limitation

Cells respond to environmental stress conditions by adjusting gene expression and protein synthesis, a role mainly filled by protein kinases signaling pathways. However, the key molecular players involved in AA-sensing regulation in other organisms are absent from the genome of malaria parasite. To identify candidate regulators, we examined a panel of 15 eukaryotic protein kinases (ePKs) which were previously shown to be non-essential for asexual erythrocytic stages of *P. berghei* development in mice^37^. By *in vitro* culturing *P. berghei* lines with loss-of-function mutations in protein kinase genes^37^ in AA depleted media (**Figure 3, Figure 3 - Figure supplement 1a,b)** we have identified nek4, eIK1 and eIK2 mutants as unresponsive to both MetD and IleD – suggesting a role for these kinases in AA sensing, albeit with some relevant particularities at the phenotypic level (**Figure 3).**

**Figure 3.**
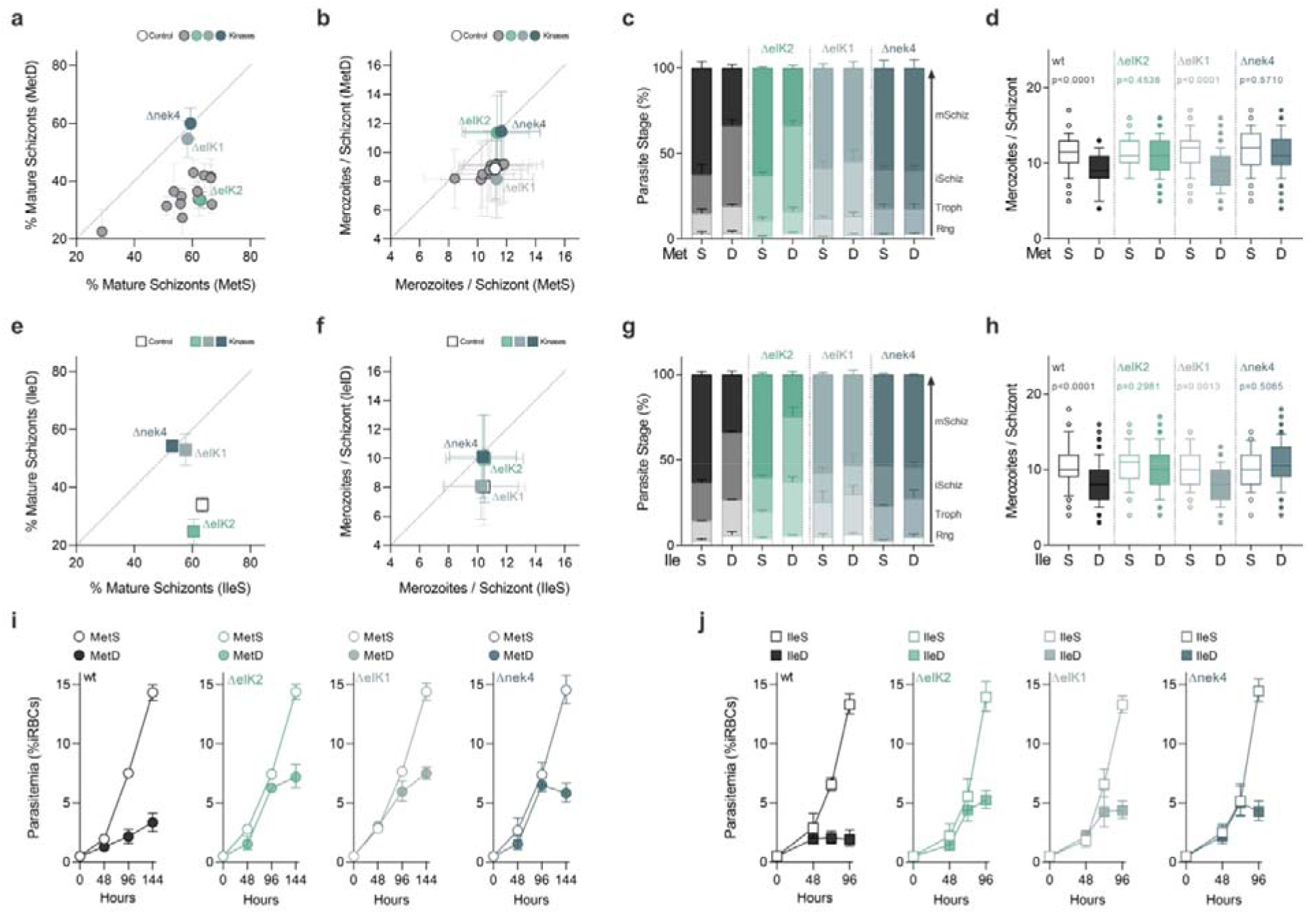
A signaling pathway comprising 3 distinct parasite kinases regulates the stress response to AA depletion. **(a, b)** Screen of 15 non-essential *P. berghei* kinases in loss-of-function mutants using the *in vitro* maturation assay. **(a)** Percentage of mature segmented schizonts and **(b)** merozoite number per schizont in wild-type and kinase knockout lines (Δnek4: PBANKA_061670; ΔeIK1: PBANKA_1308400; ΔeIK2: PBANKA_0205800) cultured in MetD-media. **(c, d)** Quantification of *P. berghei* **(c)** intra-erythrocytic developmental stages and **(d)** box plot of mean merozoite numbers of *P. berghe*i wild-type, ΔeIK2, ΔeIK1 and Δnek4 parasites cultured in MetD media. **(e)** Percentage of mature segmented schizonts and **(f)** mean merozoite numbers in wild-type and kinase knockout parasite lines, cultured for 24h in IleS- or IleD-media. **(g, h)** Quantification of **(g)** *P. berghei* intra-erythrocytic developmental stages or **(h)** box plot of mean merozoite numbers of *P. berghei* wild-type and kinase mutant ΔeIK2, ΔeIK1 and Δnek4 parasites cultured in IleD conditions. **(i, j)** Parasitemia of synchronized *P. falciparum* 3D7 WT, ΔeIK2 (PF3D7_0107600), ΔeIK1 (PF3D7_1444500) and Δnek4 (PF3D7_0719200) parasites cultured in **(i)** MetS- or MetD-media (*n*=6/ condition) and **(j)** IleS- or IleD-media (IleS, *n*=4/condition). **a, b, e, f.** All kinase knockout mutants were analyzed in triplicate and averaged, with each data point representing the **(a, e)** mean percentage of mature schizonts or **(b, f)** the mean merozoite number produced per schizont. Error bars represent the SE Diff or SD between replicates, respectively. 3 independent experiments are represented for wild-type, ΔeIK2, ΔeIK1 and Δnek4 or *N*=2 for the other knockout lines. **c, g**. Parasite staging was performed in triplicate, averaged and repeated in 2-3 independent experiments. Data is shown as mean percentage of parasites at each developmental stage, with error bars representing SEM. **d, h.** Data is represented as box-whisker plot of mean merozoite number per schizont ± SD (2-way ANOVA), with the median represented at the center line. Boxplots show the data of 2-3 independent experiments and for the following number of schizonts, wild-type: MetS, *n*=148; MetD, *n*=148; IleS, *n*=114; IleD, *n*=133; ΔeIK2: MetS, *n*=85; MetD, *n*=158; IleS, *n*=70; IleD, *n*=68; ΔeIK1: MetS, *n*=220; MetD, *n*=238; IleS, *n*=75; IleD, *n*=94 and Δnek4: MetS, *n*=106; MetD, *n*=106; IleS, *n*=90; IleD, *n*= 102. **i,j.** Data represents the mean percentage of iRBCs ± SEM (2-way ANOVA), determined in 2 independent experiments, each performed at least in triplicate. **Figure 3 – Source data 1** Phenotypic analysis of kinase mutant knockout parasites growing in MetD conditions related to Figure 3a,b. **Figure 3 – Source data 2** *In vitro* maturation and replication rates of *P. berghei* ΔeIK1, ΔeIK2 and Δnek4 parasites growing in MetD conditions related to Figure 3c,d. **Figure 3 – Source data 3** Phenotypic analysis of *P. berghei* ΔeIK1, ΔeIK2 and Δnek4 parasites growing in IleD conditions related to Figure 3e,f. **Figure 3 – Source data 4** *In vitro* maturation and replication rates of *P. berghei* ΔeIK1, ΔeIK2 and Δnek4 parasites growing in IleD conditions related to Figure 3g,h. **Figure 3 – Source data 5** Comparison of *P. falciparum* ΔeIK1, ΔeIK2 and Δnek4 kinase mutants growth in AA depleted media related to Figure 2i,j.

In MetD or IleD conditions, Δnek4 parasites mature and replicate similarly to wild-type parasites in MetS **(Figure 3a-d**) or IleS **(Figure 3e-h**). In contrast, while ΔeIK1 parasites in MetD or IleD conditions achieve maturation rates comparable to wild-type parasites in control media, the merozoite numbers are still markedly diminished **(Figure 3a-h**). Interestingly, ΔeIK2 parasites in MetD or IleD conditions exhibit the opposite response, with maturation rates markedly diminished yet, the merozoite numbers generated per schizont resemble wild-type parasites growing in control conditions **(Figure 3a-h**). A similar parasite response to MetD or IleD conditions was observed for *P. falciparum* kinase mutant knockout lines - ΔeIK1^21^, ΔeIk2^38^ and Δnek4^39^ - when assessing parasite growth **(Fig. 3i, j)** as well as intra-erythrocytic developmental and replication rates **(Figure 4 - Figure supplement 1).** Interestingly, such inability of kinase mutant knockout parasites to modulate growth – either development (ΔeIK1), replication (ΔeIK2) or both (Δnek4) – in response to AA depletion likely culminates in parasite death after some rounds of replication, as evidenced by a stalling (ΔeIK1/2) or even reduction (Δnek4) in parasitemia **(Figure 3i, j)**. The involvement of nek4, eIK1 and eIK2 in parasite adaptation to AA deficiency strongly suggests that these kinases regulate an active parasite response to AA starvation, whose modulation occurs in a stage-specific and temporally-regulated manner, a mechanism that was so far elusive. Moreover, our data strongly suggests that nek4 acts upstream to eIK1 and eIK2 in the pathway, regulating both replication - via eIK2 – or development - via eIK1.

## Parasite response to low AA implies eIF2α phosphorylation at different stages of the cell cycle

*P. falciparum* parasites growing in AA-free media exhibit increased levels of phosphorylation of the eukaryotic initiation factor 2α (eIF2α)^21^, which in eukaryotic model organisms is a well characterized mechanism that blocks translation under AA depletion^40–42^. From yeast^43^ to mammals^44^, such mechanism is coordinated by the eIF2a kinase GCN2, which detects the increased accumulation of uncharged tRNAs induced upon AA depletion^41,42^. In *Plasmodium*, three eIF2α have been identified: eIK1, eIK2 and PK4^45^ whose only described function is to phosphorylate the *Plasmodium* eIF2α orthologue under different stresses^28^. Since our screen data revealed a key role for eIK1 and eIK2 – two, of the three eIF2α kinases - in parasite response to AA depletion, we next sought to establish whether these kinases were coordinating an adaptive response mechanism, thereby modulating phosphorylation of its downstream target eIF2α.

Immunoblotting analysis of synchronized *P. falciparum* WT parasites reveals that under control conditions (MetS, IleS conditions) eIF2α phosphorylation is low at the ring stage, increasing later during schizont development (**Fig. 4**), which has been taken to be indicative of decreased protein synthesis and successful maturation^46^. By contrast, MetD- and IleD-conditions impact on eIF2α phosphorylation status (**Fig. 4**), suggestive of an early and active parasite response to low AA levels, yet with some particularities - as previously observed at the phenotypic level (**Figure 1).**

**Figure 4.**
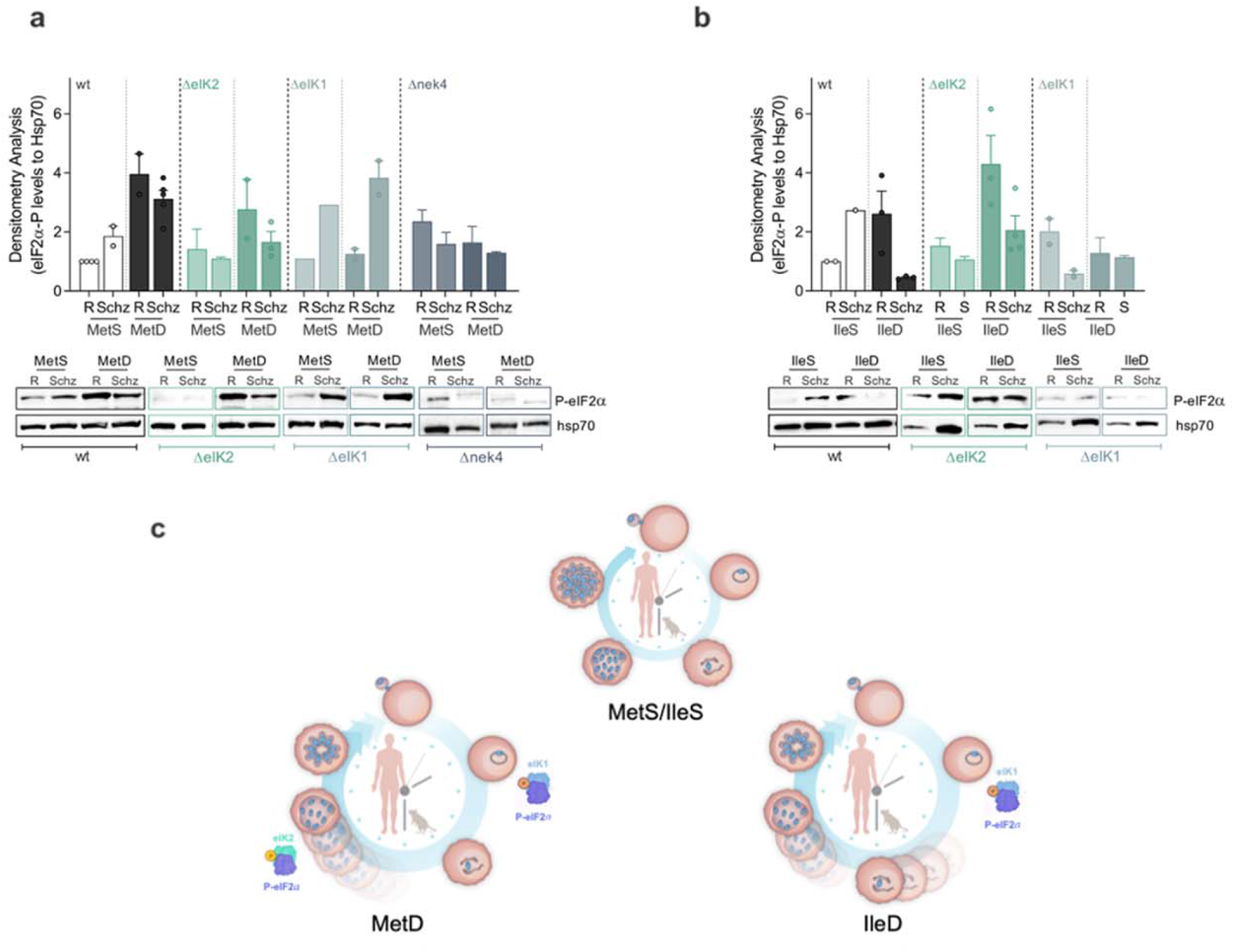
Parasite response to AA depletion results in enhanced phosphorylation of the *P. falciparum* eIF2a ortholog. **(A, B)** Time course analysis of eIF2α phosphorylation status upon AA starvation. *Pf*3D7 WT, eIK2, eIK1 and nek4 kinase knockout mutants were sorbitol-synchronized twice to achieve a ring-stage window of ± 8h interval and cultured in complete MCM medium for one developmental cycle. Parasites were cultured in **(A)** MetS- and MetD-media or **(B)** IleS- and IleD-media. Starvation pulses were performed for 6 hours at the ring-(R) or schizont-(Schiz) stage. Infected red blood cells were isolated by saponin lysis, parasite proteins were extracted, processed for western blot and probed for the phosphorylation of the translation initiation factor 2α (P-eIF2α) at the serine residue 51. Representative phospho-western blots and bar graphs quantifying phosphorylation levels of eIF2α, in ring- and schizont-stage parasites, relative to levels in ring-stage parasites growing under control conditions (MetS or IleS, respectively). Values represent the mean of eIF2α-P levels relative to hsp70 levels ± SEM, examined in 2 independent experiments. **(C)** Schematic representation of parasite’s adaptive response mechanism to AA depletion during intraerythrocytic development. *Plasmodium* parasites sense and actively respond to fluctuations in the availability of two distinct AAs – Ile and Met. Such response is heterogeneous as it depends on the AA that is depleted yet, it implies a common pathway comprising the phosphorylation of eIF2α by three distinct kinases, nek4, eIK1 and eIK2, the last two being temporally regulated at distinct stages of the life cycle, allowing parasites to fine-tune replication and development accordingly. This response results in lower replication, delayed development and ultimately, extended survival. **Figure 4 – Source data 1** Comparison of P-eIF2α levels in *P. falciparum* wt and kinase mutant knockout lines (eIK1, eIK2 and nek4) growing in MetS or MetD media related to Figure 4a. **Figure 4 – Source data 2** Comparison of P-eIF2α levels in P. *falciparum* wt and kinase mutant knockout lines (eIK1, eIK2 and nek4) growing in IleS or IleD media related to Figure 4b.

Both MetD and IleD conditions strongly induce phosphorylation of eIF2α earlier in development, at the ring stage. However, while in MetD conditions enhanced phosphorylation is sustained throughout schizont development **(Figure 4a)**, in IleD conditions eIF2α phosphorylation is quickly lost and restored to basal levels as schizont development progresses **(Fig. 4b)**. This is in agreement with our previous findings showing that IleD mainly impacts early developmental parasite stages, whereas MetD has the strongest impact during schizont development **(Figure 1 - Figure supplement 1b,c).**

Our results show that ΔeIK1 parasites are solely capable of responding to AA depletion during the schizont stage as evidenced by increased eIF2α phosphorylation levels at this particular stage, with evident loss of eIF2α phosphorylation during ring-stage development **(Figure 4a, b)**. By contrast, while ΔeIK2 schizonts are incapable of phosphorylating eIF2α, ring-stage parasites successfully respond and adapt to AA scarcity. Interestingly, Δnek4 parasites are totally unable of responding to MetD, exhibiting minimal eIF2α phosphorylation levels both at the ring- and schizont-stages **(Figure 4a).** Therefore, while eIK1 seems critical for parasite response during early-stages of development, eIK2 activity emerges as only relevant towards late stages of development - at least in MetD **(Fig. 4a).** Remarkably, nek4 activity appears as key for parasite response to AA depletion throughout the entire intra-erythrocytic development – a kinase only known to function during gametocytogenesis^47^ **(Figure 4a).**

While it has been previously shown that ring-stage parasites lacking eIK1 do not phosphorylate eIF2α in response to AA depletion, the ability of schizonts to actively respond to AA fluctuations as well as the role of eIK2 - so far only described to have a function in the sporozoite stage^21^ - and nek4 in response to AA depletion was still elusive.

Overall, the data show that *Plasmodium* parasites are capable of sensing AA fluctuations and mount active responses to individual AAs. This occurs via a common mechanism comprising the phosphorylation of eIF2α by two different kinases - eIK1 and eIK2 – whose function is likely coordinated by nek4 in a temporally-regulated manner, with each kinase affecting a specific, but sequential, stage of development.

## Discussion

Nutrient scarcity is an important selective pressure that has shaped the evolution of many organisms and most cellular processes. While nutrient sensing pathways have been established and studied in model systems, the capacity of malaria parasites to sense nutrients has only recently started to be revealed. This arises from the fact that *Plasmodium* parasites do not seem to possess any of the canonical eukaryotic nutrient-sensing pathways, including SNF/AMPK, TOR and GCN2 biosynthetic pathways. Notably, while the canonical SNF/AMPK energy-sensing pathway was thought to be absent, we have recently shown that a parasite kinase, KIN, is a putative functional AMPK homologue, working as a broad metabolic/energy sensor that drives an active and coordinated parasite response to energy fluctuations^12^.

On the other hand, the TOR complex - the master regulator of cell growth in eukaryotes - is completely absent in *Plasmodium*, which has been shown to possess only a rudimentary AA starvation-sensing eukaryotic initiation factor 2α (eIF2α), that does not directly promote parasite survival upon AA starvation^21,24^. In many model organisms, it has been shown that GCN2 senses the uncharged tRNAs that accumulate upon AA deprivation. GCN2 attenuates translation, which not only consumes AA, but is also one of the most energy-demanding cellular processes^16^. We now clearly show that asexual blood stage parasites are able to actively adapt to a reduction in AAs through 3 distinct kinases: nek4 – a kinase so far described to function during gametocyte development^47^ – eIK1 and eIK2. Additionally, our data suggests that these two distinct eIF2α kinases function in a temporally-regulated manner, with eIK1 regulating developmental stage transition but not replication and eIK2 showing the opposite behavior, as it regulates replication but not development. This translates in a differential dynamics of P-eIF2α observed for each kinase mutant throughout development under AA-deprived conditions. Indeed, our data suggests that in order to survive under AA deficiency, malaria parasites must be able to modulate replication levels as well as cell cycle pace – an adaptive mechanism that comprises phosphorylation of eIF2α. Thus, the decreased replicative capacity and reduced cell cycle pace observed in wt parasites translates an ability to successfully respond to AA depletion – a response that relies on the enhanced phosphorylation of eIF2α, which must be sustained throughout development (ring- and schizont-stage). Albeit at the cost of the progeny generated, such adaptive response ensures that parasites can commensurate nutrient availability with growth, ultimately extending survival. By contrast, the inability of Δnek4 parasites to control replication and growth under AA deficiency, which closely relates with the complete loss of eIF2α phosphorylation, first translates in high growth rates yet, it rapidly culminates in parasite death. Interestingly, ΔeIK1 and ΔeIK2 parasites exhibit an intermediate phenotype under AA deficiency, with each kinase regulating either replication or growth and phosphorylating eIF2α in a stage-specific manner. Although ΔeIK2 ring-stage parasites are able to adapt, schizonts fail to phosphorylate eIF2a in response to AA deficiency, which indicates that parasite replicative rates might be pre-determined at this stage. By contrast, while ΔeIK1 schizonts are able to adapt, ring-stages fail to phosphorylate eIF2a in response to AA deficiency, which corroborates the ability of this mutants to successfully complete the cell cycle and develop into mature schizonts. Indeed, eIF2α phosphorylation at the schizont stage, under basal conditions, has been shown to be key for the successful maturation of schizonts further corroborating our data^28^.

Interestingly, such stage-specific and sequential activation of eIF2α kinases constitutes a major hallmark of the integrated stress response in other eukaryotes^48^ including the closely related apicomplexan parasite *Toxoplasma gondii*^49^. Furthermore, although adaptation to low AA levels entails a common AA-sensing pathway, comprising nek4 and the phosphorylation of eIF2α by eIK1 or eIK2, the overall response observed for the two AAs tested, Met and Ile, exhibits some particularities, in such a way that each AA differentially impacts on the cell cycle phenotypes observed, as well as in the dynamics of eIF2α phosphorylation. This is not surprising when taking into account current evidence in other eukaryotes where the GCN2-mediated response to AA starvation differs according to the specific AA that is depleted^50–53^ - with Met starvation leading to the most striking decrease in protein translation in mammalian cells^54^.

Still to be disclosed is whether nek4 acts upstream of eIK1 and eIK2, which would support our phenotypic analysis showing a role for nek4 in both replication and development. Additional studies addressing whether under AA depleting conditions, eIK1 and eIK2 activate nek4 in asexual blood stages would provide more insight on how this AA sensing pathway is regulated in the parasite. Additional studies addressing the role of PK4 – already shown to phosphorylate eIF2α during schizont development^27,28^ - in parasite response to AA starvation may also provide additional insights into the dynamics eIF2α phosphorylation as well as the function of these kinases as critical components of a GCN2-like sensing pathway in the parasite. Moreover, future experiments addressing the phosphorylation status and activity of *Maf1*, the sole downstream component of the TOR pathway identified to date in *Plasmodium*, under AA deficiency may disclose whether a TOR pathway operates in *Plasmodium* and on how it cross-talks with the GCN2 pathway to ensure a suitable cellular adaptation to nutritional fluctuations

While malaria eradication is highly desirable, its accomplishment seems to be rather difficult. This relies primarily on the emergence of parasite resistance to artemisinin-based combination therapies (ACTs), the frontline treatment against *Plasmodium falciparum* malaria. While the mechanism of action of artemisinin (ART) is not fully understood, it has been shown that drug treatment blocks the uptake of host cell hemoglobin while markedly increasing general protein damage, resulting in delayed parasite clearance and the persistence of hemoglobin non-degrading ring-stage forms. Therefore, ART treatment is mostly effective at later stages of development when parasites exhibit high metabolic activity^55–57^. Moreover, besides inducing a transient reduction in cell cycle pace, ART treatment also enhances eIF2α phosphorylation, resembling amino acid withdrawal-induced phenotypes. Importantly, increased eIF2α phosphorylation has been shown to be essential for ART-induced latency and parasite recrudescence^58^. Such a programmed state of low metabolic activity, both upon ART-treatment or AA withdrawal - could render parasites insensitive to a broad range of inhibitors, as already shown for the exoerythrocytic stages of other human malaria parasites^59^. As such, by unveiling and blocking the molecular players involved in *Plasmodium* AA sensing pathways – namely nek4 or the two eIF2α kinases, eIK1 and eIK2 – represents a novel and promising strategy to prevent parasite ability to transition into this latent phase, hindering thereby malaria recrudescence following ART treatment. Additionally, interventions targeting these kinases would transform highly replicating, and as such virulent parasites into attenuated parasites leading to low parasitemia, previously shown to be protective in humans^60^.

## Methods

### Chemicals and Reagents

Roswell Park Memorial Institute (RPMI) 1640 medium, no glutamine (21870076), HEPES (15630), Gentamicin (15750), L-glutamine (25030-024), Albumax II (11021-029) and SYBR green I (S7567) were purchased from Gibco, Thermo Fisher Scientific. RPMI 1640 Medium, no isoleucine (R9014) was purchased from US Biologicals. Bovine Serum Albumin (BSA) Fraction V (MB046) was purchased from NZYTech. RPMI 1640 no methionine, no cysteine (R7513), Glucose (G6152), Saponin (47036), Spermine (S4264), Spermidine (S0266), Putrescine (D13208), L-Glutathione reduced (G4251), L-Homocysteine (69453) and Trimethoprim (TMP) were purchased from Sigma-Aldrich. S-Adenosylmethione-1,4-butanedisulfonate (NA58198) was purchased from Biosynth Carbosynth. Paraformaldehyde was purchased from Santa Cruz Biotechnology.

### Mice, diets and treatments

Male C57BL/6J and BALB/c wild-type mice, aged 5 to 8 weeks, were purchased from Charles River Laboratories (Saint-Germain-sur-l’Arbresle, France). All mice used in this study were housed in the Rodent Facility of Instituto de Medicina Molecular João Lobo Antunes (Lisboa, Portugal), five per cage and kept in specific-pathogen-free conditions. Mice were randomly assigned to different experimental conditions and allowed free access to water and food. Blinding was not possible in MetD-fed mice as mice exhibit a clear difference in body weight. All experimental procedures involving mice were reviewed by iMM’s Animal Welfare Body (ORBEA-iMM), licensed by the national regulator, Direcção Geral de Alimentação e Veterinária (DGAV), and performed in strict compliance with national and European regulations. Methionine-sufficient (MetS) and deficient (MetD) diets were manufactured and purchased from SSniff (Soest, Germany). TMP was provided in drinking water at 0.25mg per mL starting two days prior to infection and changed every 48 hours.

### Parasite lines

The following parasite lines were employed in this study: GFP-expressing *P. berghei* ANKA (GFPcon, clone 259cl2), obtained from the Leiden Malaria Research Group; *P. berghei* ANKA kinase deficient parasite lines, were previously generated^37^; *P. falciparum* NF54 and *P. falciparum* 3D7 were obtained through MR4 (www.mr4.org); *P. falciparum sams*-glmS transgenic parasite line was kindly provided by Björn Kafsack (Weill Cornell Medical College, New York, USA) and the *P. falciparum* 3D7 kinase deficient parasite lines were kindly provided by Mathieu Brochet (University of Geneva, Geneva, Switzerland) and Christian Doerig (Monash Biomedicine Discovery Institute, Clayton, Australia).

The *P. berghei* SAMS-DD parasite line was generated by double crossover homologous recombination of the p*Pbsams*.DD.HA construct at the *sams locus* in a GFP expressing parental line (line-507cl1)^61^. The p*Pbsams*.DD.HA construct contains a truncated *Pbsams* ORF in fusion with a destabilizing domain (DD) – a mutant form of the *E. coli* dihydrofolate reductase (ecDHFR) protein, engineered to be degraded^30^ - an HA tag and also a cassette for transgenic expression of the human *dhfr* - conferring resistance to pyrimethamine-which is flanked by the *sams* 3’UTR (**Figure 1 - Figure supplement 1d**). Transfection was performed using a protocol described elsewhere^61^. Briefly, blood from a BALB/c mouse infected with the parental line was collected and cultured for 16 hours *in vitro*. Mature schizonts were purified by a Nycodenz gradient and transfected using Amaxa electroporation system (Lonza). Transfected merozoites were injected into the tail vein of a BALB/c mouse, positively selected by providing pyrimethamine in drinking water (70μg/ml) and harvested on day 7-10 post transfection. Pyrimethamine resistant, transgenic parasites were dilution cloned before further experimentation and genotyped by PCR (**Figure 1 - Figure supplement 1e**) using the following DNA oligonucleotide sequences as primers:

WT locus: p1 (fw): TAGGTACCGAGGAAATTTTCTATTTACTTCG p3 (rev): ATGCGGCCGCCAACTAATAAAATCCAGGAAATA
Modified locus: p1 (fw): TAGGTACCGAGGAAATTTTCTATTTACTTCG p4 (rev): ATGCGGCCGCTCATCGCCGCTCCAGAATCTC
QC: p1 (fw): TAGGTACCGAGGAAATTTTCTATTTACTTCG p2 (rev): TAGGGCCCATTTTTTAAAACATTTTTTTCGTG

### Mice infections and *in vivo* Parasitemia Determination

Infection of experimental mice was performed by intraperitoneal (i.p.) inoculation of 1×10^6^ infected RBCs (iRBCs) into C57BL/6J or BALB/c mice, obtained by passage in the correspondent genetic background mice. For GFP-expressing parasites, peripheral blood parasitemia was determined by flow cytometry analysis of one drop of tail blood collected in 200 μL PBS. A total number of 1 million events (day 1-2 of infection) or 100-200 thousand events (onwards) were acquired on a BD LSR Fortessa using the BD FACSDiva^™^ Software (v6.2) and the parasitemia expressed as the percentage of GFP^+^ RBCs. For non-GFP expressing parasite lines, parasitemia was monitored by Giemsa-stained thin blood smears^62^. A total of 5-10 thousand RBCs per slide, was randomly acquired and the percentage of infected RBCs (iRBCs) – parasitemia – was semi-automatically counted using Fiji software (version 2.1.0) (https://imagej.net/software/fiji/).

### *P. falciparum* culture and Parasitemia Determination by flow cytometry analysis

*P. falciparum* NF54, 3D7 and derived knockout clones were maintained in culture^63^ at 5% hematocrit in human O+ erythrocytes (Instituto Português do Sangue e Transplantação). Of note, whereas the *P. falciparum* NF54 parasite line exhibits a 48h cell cycle length, the *P. falciparum* 3D7 line shows an accelerated asexual blood stage cycle of 36-42h. Parasites were cultured in Malaria Complete parasite Media (MCM) prepared by supplementing RPMI 1640 no glutamine, with 2mM L-glutamine, 2g/L NaHCO_3_, 5mM glucose, 25mM Hepes, 100μM hypoxanthine, 10 μg/mL gentamicin, and 10% Albumax II - except for the AA starvation assays. For such experiments, MCM media was prepared by adding all compounds to RMPI lacking either methionine (MetD) or isoleucine (IleD). Respective control media, containing methionine (MetS) or isoleucine (IleS) were prepared by supplementing deficient media with the appropriate AAs, at the concentrations found in regular RPMI 1640 (100μM Met, 382μM Ile). Synchronous *P. falciparum* cultures were obtained according to previously described protocols^62,64^. To achieve synchronization, *P. falciparum* parasites were treated twice with 5% sorbitol to achieve a ring-stage window of ± 8h interval in the previous cycle. To do so, cultures were allowed to grow up to 5-10% parasitemia and harvested when parasites were mostly at the ring stage. Parasite pellets were treated with 5% sorbitol for 10 minutes at RT, centrifuged at 600x g and washed thrice in MCM. Parasitemia was assessed as well as parasite morphology and parasites were subcultured at 5% hematocrit for one additional cycle. The sorbitol treatment was repeated after one cycle – approximately 48h for NF54 and 40h for 3D7 – as previously described^64^. Ring-stages were then cultured at 2% hematocrit and the starting parasitemia adjusted to 0.5% in 96 well plates containing 200μl of the respective media. Parasitemia in each developmental cycle was analyzed by flow cytometry analysis after SYBR Green staining^65^. Briefly, 50 μL of *P. falciparum* cultured samples were mixed with the same volume of SYBR Green I solution, previously diluted in water (1:1000). Samples were incubated in the dark for 20 min at room temperature and washed in 500 μL PBS. Samples were acquired on a BD LSR Fortessa using the BD FACSDiva™ Software (v6.2). A total of 100,000-200,000 events was analyzed per condition. The data was further analyzed on FlowJo X (v10.7.1, TreeStar) using the gating strategy detailed in **Figure 1 - Figure supplement 2a-c.**

### *P. berghei ex vivo* Maturation Assays

*P. berghei* blood-stage parasites were collected by cardiac puncture of BALB/c-infected mice (1-2% parasitemia), washed in pre-warmed incomplete RPMI and cultured in 96-well plates containing complete RPMI culture medium, prepared as follows: RPMI 1640, no glutamine (21870076), supplemented with 5mM glucose, 2mM L-glutamine, 25mM HEPES, 2g/L NaHCO_3_, 50μg/L Gentamicin and 20% of Fetal Bovine Sera (FBS). Homemade complete medium sufficient for methionine (MetS) was prepared by supplementing RPMI 1640 no methionine, no cysteine (R7513) with the respective amino acids at the concentration found in regular RPMI (208μM and 100μM, respectively); whereas methionine deficient media (MetD) consisted of RPMI 1640 no methionine, no cysteine (R7513) medium supplemented only with cysteine. Isoleucine-sufficient (IleS) media was prepared by supplementing RPMI 1640 Medium, no isoleucine (IleD; R9014) with the concentration of the respective amino acid found in complete RPMI media (382 μM). Ninety-six-well plates were incubated at 37°C in an atmosphere of 5% O_2_, 5% CO_2_ and 90% N_2_ for 18-30h, allowing parasites to mature from rings/young trophozoites into segmented schizonts.

### *In vitro* quantification of parasite stage progression and merozoite number

Parasite cell cycle progression was assessed at distinct time points - 18h (*data not shown*), 24h and 30h for the rodent malaria parasite - or every 12h for the human malaria parasite. Parasite staging was performed by microscopic analysis of 1-10 thousand, randomly acquired, Giemsa-stained RBCs. Mature schizonts were assessed by microscopic analysis of Giemsa-stained smears and manually quantified using ImageJ software. Fifty to one hundred segmented schizonts with clearly individualized merozoites containing a single hemozoin crystal were quantified per condition. Additionally, rodent blood-stage nuclei were stained with DAPI and merozoite number was determined by fluorescence microscopy.

### *Ex vivo* Screen of Met-downstream metabolites

To screen for methionine-downstream metabolites, wild-type parasites were cultured in MetS or MetD media and IleS or IleD media and further supplemented with each metabolite - and the appropriate vehicle (as control). The final concentration used for each metabolite was set according to the physiological ranges described during a malaria infection or as previously described elsewhere, as follows: 1.5mM S-adenosylmethionine (SAM)^66^, 1mM glutathione (GSH)^67–69^, 20μM homocysteine (Hcy)^70^, 1mM putrescine (Putr)^71^, 1mM spermine (SpM)^66^, and 1mM spermidine (SpD)^66^. For *P. berghei*, an *in vitro* maturation assay was employed with each metabolite being added to the media and incubated for 24- or 30 hours. For *P. falciparum*, each metabolite was added to the culture media at day 4 of *in vitro* culture and incubated throughout one complete intra-erythrocytic developmental cycle. Ninety-six-well plates were incubated at 37°C in an atmosphere of 5% O_2_, 5% CO_2_ and 90% N_2_. The effect of metabolite supplementation on parasite stage progression and schizont development was assessed by microscopy of Giemsa-stained smears as described above.

### Thin Layer Chromatography Determination of Polyamines

Polyamines were separated by thin-layer chromatography as previously described^72,73^. For all samples, cells were collected and centrifuged. Pellets were washed with Phosphate Buffered Saline (PBS) and then resuspended in 100μL of 2% perchloric acid. Samples were then incubated overnight at 4°C. One hundred μL of supernatant was combined with 200μL of 5 mg/ml dansyl chloride (Sigma Aldrich) in acetone and 100 μL of saturated sodium bicarbonate. Samples were incubated in the dark overnight at room temperature (RT). Excess dansyl chloride was cleared by incubating the reaction with 100μL 150 mg/mL proline (Sigma Aldrich). Dansylated polyamines were extracted with 50μL toluene (Sigma Aldrich) and centrifuged. Five μL of sample was added in small spots to the TLC plate (silica gel matrix; Sigma Aldrich) and exposed to ascending chromatography with 1:1 cyclohexane: ethylacetate. Plate was dried and visualized via exposure to UV.

### Amino acid starvation assays for SDS-PAGE analysis

*P. falciparum* 3D7 parasites and knockout parasite lines, namely the two eIF2α kinases^45^, *Pf*ΔeIK1 (PF3D7_1444500) – previously described to closely cluster with mammalian GCN2 - and *Pf*ΔeIK2 (PF3D7_0107600) - were maintained as described above in MCM media supplemented with human O+ erythrocytes at 2% hematocrit. Parasites were sorbitol-synchronized to the ring-stage and allowed to grow in culture for up to 5% parasitemia. Parasites were collected at two distinct stages of the intra-erythrocytic developmental cycle: ring (12h) or schizont (38h). Ring- or schizont-stages were pelleted by centrifugation, washed twice in PBS, pH7.5, equally partitioned and cultured at 37°C with 5% CO_2_ for a 6-h pulse in media deficient for methionine- or isoleucine-(MetD or IleD) and the respective control media (MetS or IleS) - prepared as previously described. Cultures were then harvested by centrifugation at 1300x g for 5 minutes at RT and further processed for parasite pellet isolation.

### Parasite pellet harvesting and isolation for SDS-PAGE

Pelleted erythrocytes were washed in equal volume of PBS, harvested by centrifugation at 1300x g for 2 minutes at 4°C and further lysed on ice by treatment with 0.15% saponin in PBS. Parasite pellets were isolated by centrifugation at 5500x g for 10min, 4°C and parasites were washed twice in PBS. Blood-stage parasite proteins were isolated by lysis in 50-200μL of radioimmunoprecipitation assay (RIPA) buffer and washed in PBS with cOmplete^™^ protease inhibitor cocktail (Roche). Samples were resuspended in 5x SDS-Laemmli buffer and denatured at 95°C.

### Immunoblotting Analysis

Denatured parasite proteins were resolved on 10% SDS-polyacrylamide gel and transferred to 0.2 μm nitrocellulose membranes (Bio-Rad) using wet-transfer in Towbin buffer (25 mM Tris, 192 mM Glycine (Sigma) and 20% ethanol in ddH2O). Membranes were blocked in 5% BSA in PBS-0.1% Tween 20 (PBST) at RT for 1h, with rocking. Membranes were then incubated with the appropriate primary antibodies, diluted 1:1000 in 5% BSA in PBST, for overnight at 4°C. Membranes were washed thrice in 0.1% PBST for 10 minutes (RT) followed by incubation with appropriate secondary antibodies (conjugated to Horseradish peroxidase) diluted 1:10 000 in 5% BSA in 0.1% PBST, for 60 min at RT. Membranes were washed thrice and bound antibodies were detected by using Immobilon ECL Ultra Western HRP Substrate (Merck Millipore) on the ChemiDoc XRS+ system.

The primary antibodies used for probing membranes were rabbit α HA (clone C29F4, Cell Signaling Technology, Danvers, MA, USA), rabbit α phospho-eIF2α (S51) (119A11, Cell Signaling Technology) and mouse α *Pb*hsp70 (produced in house). The secondary antibodies used were: goat α mouse IgG, HRP conjugate (BML-SA204-0100, Enzo Life Sciences, Lausen, Switzerland) and goat α rabbit IgG, HRP-linked Antibody (7074, Cell Signaling Technology).

### Immunofluorescence Assay

Immunofluorescence analysis of SAMS-DD-HA protein levels was performed in blood smears of *P. berghei* parasites cultured *in vitro* for 24h. MetS-fed BALB/c mice infected with *P. berghei* SAMS-DD parasites were either treated, or not, with TMP in drinking water two days prior to infection to achieve SAMS stabilization or knockdown, respectively. Parasites were cultured for 24h as previously described for the *in vitro* maturation assays and blood smears were fixed in 4% PFA-PBS solution for 10 min at RT. Fixed parasites were washed in PBS and permeabilized in 0.1% PBS-Triton X-100 solution for 10 min at RT. Cells were then blocked in 3% BSA for 1h at RT. Primary antibodies were diluted 1:400 in blocking solution and incubated at 4°C, overnight. Slides were rinsed thrice in PBS and incubated in secondary antibodies or dyes, diluted 1:400 or 1:1000, respectively, in blocking solution. Stained samples were rinsed in PBS, mounted in Fluoromount G and allowed to dry overnight. Primary antibodies used for fluorescence microscopy include: rabbit α HA (C29F4, Cell Signaling Technology) and mouse α *Pb*hsp70 (produced in house). Secondary antibodies and dyes used for fluorescence microscopy include Alexa-488 conjugated donkey α mouse GFP (A21311, Thermo Fischer Scientific), Alexa-647 conjugated donkey α rabbit IgG (A32795, Thermo Fischer Scientific) and DAPI for nuclear staining (D1306, Invitrogen, Thermo Fischer Scientific). Slides were imaged in Zeiss LSM 710 confocal laser point-scanning fluorescence microscope (ZEN 2 software, Blue edition) and merozoite counting was performed by the use of ×63 oil objective.

### Statistical Analysis

Significance was determined by different statistical tests on the GraphPad (Prism, version 8.4.3) software, as follows: Mann-Whitney U tests was used for comparisons between two different groups or conditions, two-way ANOVA was used to compare parasitemia, development and merozoite numbers between the control counterparts and the following experimental conditions: MetD-, IleD-, kinase mutants or metabolite supplementation assays. The log-rank (Mattel-Cox) test was used to compare the survival distributions of two groups. Significance was considered for *p* values below 0.05. The outliers in the boxplots represent 10% of data points. Biological replicates *(n)* indicated in figure legends refer to the number of mice, number of schizonts, or number of culture wells. The number of independent experiments is referred to as *N*. Sample sizes were chosen on the basis of historical data. No statistical method was employed to predetermine sample size.

### Reporting summary

Further information on research design is available in the Nature Research Reporting Summary linked to this article.

## Data availability

All data generated or analysed during this study are included in this manuscript and its Supplementary Information. Source data underlying the graphs presented in the main figures and figure supplements are available. Full blots and replicates are shown in the source data.

## Supplementary information

This file contains the uncropped source data for Figure 2f, Figure 4a, Figure 4b and Figure 1 – Figure supplement 1e,f.

## Acknowledgments

From IMM-JLA, Lisbon, Portugal, we would like to acknowledge Inês Bento for discussions, critical input and reading of the manuscript, Ângelo Ferreira Chora for guidance with flow cytometry analysis and for critical reading of the manuscript, Yash G. Pandya for critically reviewing this manuscript, the Bioimaging Facility for assistance with microscopy experiments, the Rodent Facility for overall animal husbandry and support with mice handling and the Flow Cytometry facility for assistance with equipment handling. We would like to thank Mathieu Brochet (University of Geneva Department of Microbiology and Molecular Medicine, Genève, Switzerland) and Christian Doerig (Monash Biomedicine Discovery Institute, Clayton, Australia) for kindly gifting the *P. falciparum* kinase mutant parasite lines: ΔeIK1, ΔeIK2 and Δnek4. We would like also to express our gratitude to Arthur Scherf (Biology of Host-Parasite Interaction, Institut Pasteur, Paris, France) for discussions and for critically reviewing this manuscript.

Work at iMM-JLA was supported by Fundação para a Ciência e Tecnologia (PTDC/BIA-MOL/30112/2017) and the “La caixa” Banking Foundation (HR17-00264-PoEMM) grants attributed to V.Z.L. and M.M.M., respectively. Work at the Weill Cornell Medicine (BK. and CT) was funded by NIH 1R01 AI141965 (BK), NIH 1R01 AI138499 (BK), Alice Bohmfalk Charitable Trust Research Grant (B.K.), NIH 5F31AI136405-03 (C.H.). Work at the Loyola University Chicago (B.M. and V.C.) was supported by funds from NIH NIGMS R35GM138199. M.I.M., S.M. and V.Z.L. were supported by Fundação para a Ciência e Tecnologia, Portugal (PD/BD/135454/2017, DL57/2016/CP1451/CT0010 and DL 57/2016/CP1451/CT0022 respectively).

---

**Figure 1 – Figure supplement 1.**
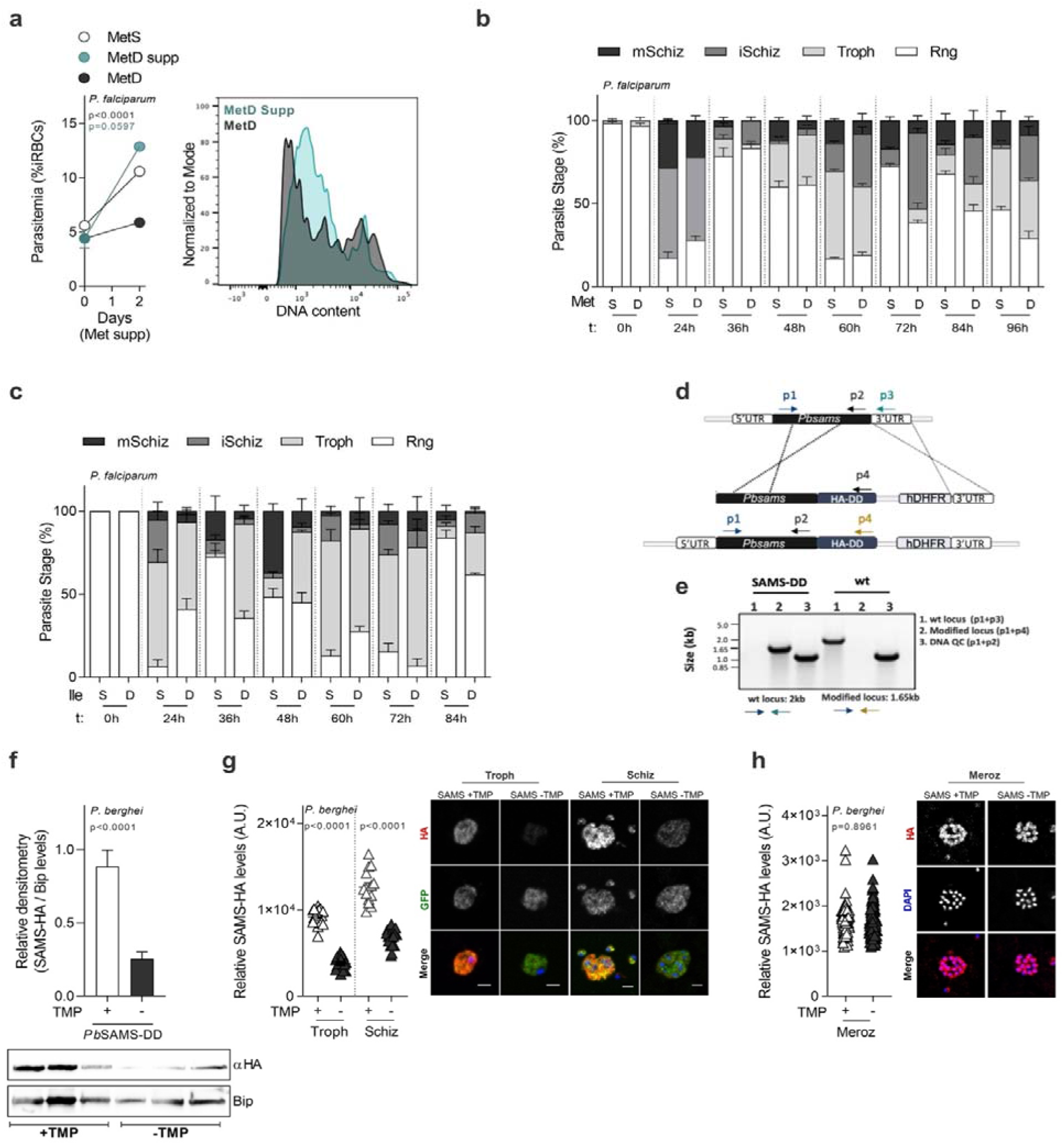
AA depletion impacts parasite intra-erythrocytic development but not viability. **(a)** Parasitemia of synchronized *P. falciparum* WT parasites cultured in MetS-, MetD- or MetD-media supplemented with methionine (MetD supp), by flow cytometry after SYBR Green staining. Methionine was provided to the media at the concentration found in RMPI (100μM) from day 4 to day 6 of *in vitro* culture. (**b, c)** Full course analysis of *Pf* 3D7 WT developmental stage progression in **(b)** MetS- or MetD-media or **(c)** IleS- or IleD-media for 2 developmental cycles. Parasite staging was performed by microscopy analysis of Giemsa-stained blood smears. **(d)** Double crossover homologous recombination at the *Pbsams* locus and the 3’UTR was employed to introduce a fusion of *Pbsams* ORF with the destabilizing domain system (DD) and an HA tag (*Pb*SAMS-DD line). SAMS stabilization was achieved by providing TMP to mice in drinking water (0.25mg/mL) and transgenic parasites were positively selected by pyrimethamine treatment (70μg/ml). Annealing sites for genotyping primers are illustrated and primer sequences are given in the methods section. **(e)** Agarose gel image, representative of 3, showing diagnostic PCR products from *P. berghei* SAMS-DD and *P. berghei* WT genomic DNA, after dilution cloning of the *Pb*SAMS-DD transgenic parasite line. **(f)** Densitometry analysis of SAMS-HA-DD protein levels in BALB/c mice infected by intraperitoneal (i.p.) inoculation of 1×10^6^ *Pb*SAMS-DD infected RBCs (iRBC). SAMS protein levels were probed for the HA-tag (*Pb*SAMS-HA-DD line; αHA, 70KDa). **(g, h)** Representative image and quantification of SAMS-HA-DD protein levels at the **(g)** trophozoite-(Troph), schizont-(Schiz) and **(h)** merozoite-stage (Meroz) of parasites developing in BALB/c mice treated or not with TMP, measured by fluorescence microscopy after probing for HA-tag. **a.** Data represents the mean percentage of iRBCs ± SEM (2-way ANOVA), determined in 2 independent experiments, each performed at least in triplicate. **b,c.** Parasite staging was performed in triplicate, averaged and repeated at least in 2-3 independent experiments. Data is shown as mean percentage of parasites at each developmental stage with error bars representing SEM. **f.** Western blot image, representative of 2, showing SAMS stabilization/destabilization after TMP treatment. Each lane represents an individual mouse. Data is shown as mean ± SEM of SAMS-HA-DD densitometry levels normalized to Bip levels (Mann-Whitney). Analysis was performed in 3-4 mice per conditions, averaged and repeated at least in 3 independent experiments. **g,h.** Scatter dot plots represent SAMS intensity levels, measured in 2 independent experiments (n=2 mice/condition). Data is shown as SAMS mean intensity values inside *Pb*SAMS-DD +TMP and *Pb*SAMS-DD -TMP parasites, normalized to parasite area (Mann-Whitney). The black line represents the mean and scatter dot plots show the data for the following number of parasites: Trophozoite-stage: SAMS-DD: +TMP, *n*=20; -TMP, *n*=20; Schizont-stage: SAMS-DD: +TMP, *n*=15; -TMP, *n*=21 and Merozoite-stage: SAMS-DD: +TMP, *n*=102; -TMP *n*=108. Scale bar = 10μm. **Figure 1 – Figure supplement 1 - Source data 1** Rescue of MetD-induced attenuated growth in *P. falciparum* parasites by Met supplementation related to Figure 1, Figure supplement 1a. **Figure 1 – Figure supplement 1 - Source data 2** Comparison of cell cycle progression in *P. falciparum* parasites growing in AA sufficient or deficient media, related to Figure 1, Figure supplement 1b,c. **Figure 1 – Figure supplement 1 - Source data 3** *P. berghei* SAMS-HA-DD protein expression levels in mice treated or not with TMP related to Figure 1, Figure supplement 1f. **Figure 1 – Figure supplement 1 - Source data 4** *P. berghei* SAMS-HA-DD protein expression levels upon TMP treatment or not, in trophozoite-, schizont- and merozoite-stages related to Figure 1, Figure supplement 1g,h.

**Figure 1 – Figure supplement 2.**
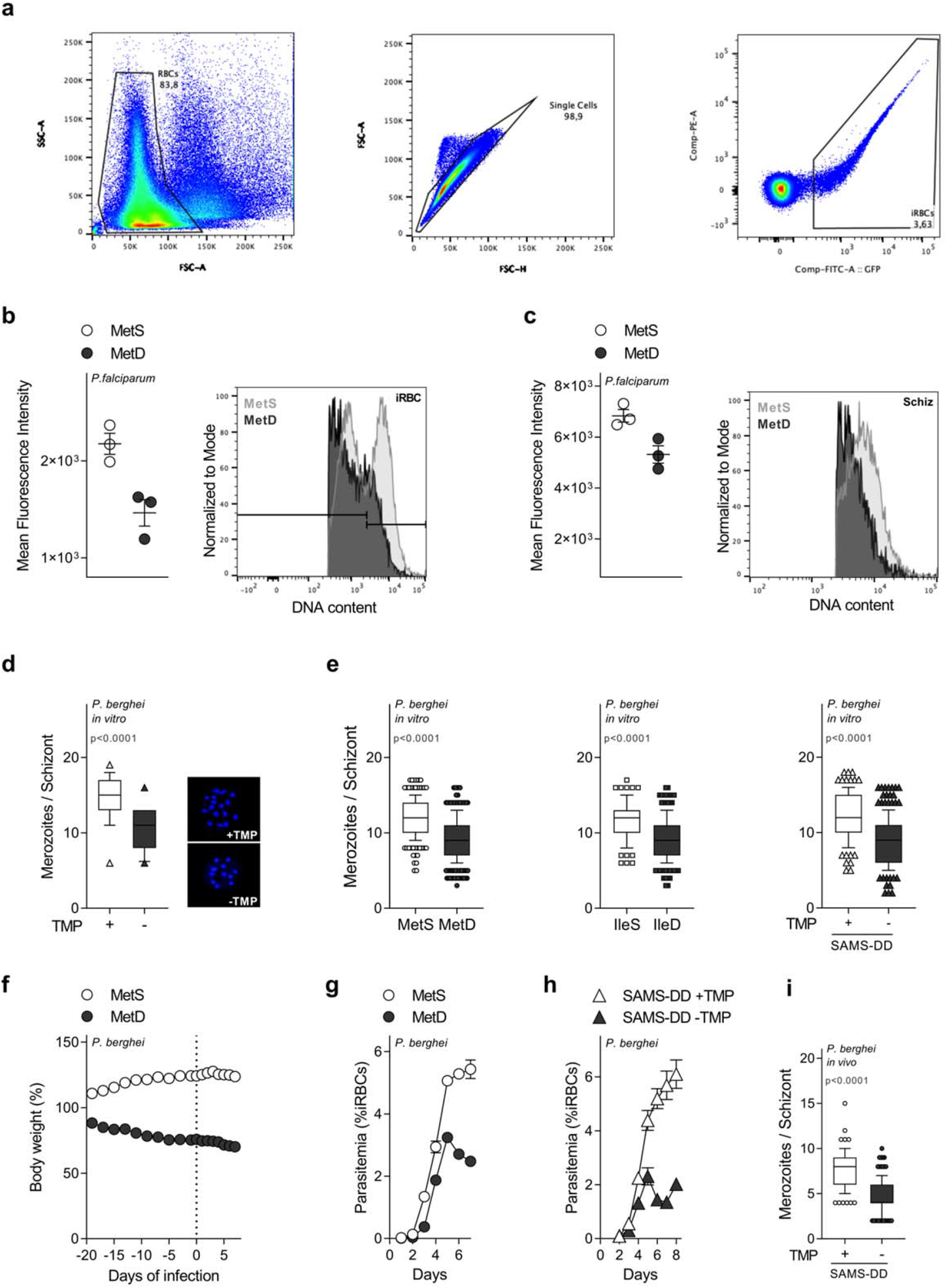
Effect of AA depletion *in vitro* and *in vivo*. **(a)** Representative image of flow cytometry plots and gating strategy for analysis of synchronized *P. falciparum* parasites cultured in MetS- or MetD-media after SYBR-Green staining. Samples were analyzed on BD LSRFortessa and the number of total acquired events ranged from 50 000 to 10 000/condition. The data was further processed on FlowJo^™^ 10 Software (Tree Star Inc.). Single RBCs were selected on the basis of their size (left and middle panel) and, subsequently, on the DNA content, excluding false positives associated to RBC autofluorescence (right panel). **(b, c)** Mean Fluorescence Intensity (MFI) of **(b)** RBCs infected with synchronized *Pf*NF54 WT parasites or **(c)** of *Pf*NF54 WT schizonts cultured as in **a.** Histogram plot showing the fluorescent intensity comparison between MetS and MetD. As shown in panel **b**, schizonts express the strongest SYBR-Green signal detected in the FITC channel. **(d)** Boxplot of mean merozoite numbers in *P. berghei* SAMS-DD parasites - treated or not with TMP, using the *in vitro* maturation assay. Mature schizonts were visualized by microscopy after DNA staining with DAPI and quantified using the ImageJ counter tools. Representative fluorescent microscopy image shows the mean merozoite number comparison between SAMS-DD +TMP and SAMS-DD -TMP parasites. **(e)** Box plot of mean merozoites number per schizont of *P. berghei* parasites after *in vitro* culture for 30h, in MetD-media or IIeD-media and in SAMS knockdown parasites. **(f)** Body weight change of BALB/c male mice under a long-term methionine restriction-regimen (MetD) or in regular diet - sufficient for Met (MetS). **(g)** Parasitemia of BALB/c mice under a MetS- or MetD-regimen, infected by intraperitoneal *(i.p.)* injection of 1×10^6^ GFP-expressing *P. berghei* ANKA WT-infected RBCs. **(h)** Parasitemia of MetS-fed BALB/c mice infected with *Pb*SAMS-DD parasites and treated, or not, with TMP in drinking water. **(i)** Box plot of mean merozoite number per schizont of *P. berghei* SAMS-DD parasites developing *in vivo* in MetS-fed BALB/c mice treated or not with TMP. **b, c.** Data is shown as scatter dot plot and represents 1 of 2 independent experiments *(n=3* mice/condition); black line represents the mean. **d,e.** Data is represented as box-whisker plot of mean merozoite number per schizont ± SD (Mann-Whitney), with the median represented at the center line. Boxplots show the data of 2-4 independent experiments and for the following number of schizonts: (d) MetS, *n*=20; MetD, *n*=20; (e) MetS, *n*=261, MetD, *n*=340; IleS, *n*=113, IleD, *n*=192; SAMS-DD +TMP, *n*=163, SAMS-DD -TMP, *n*=200. **f.** Data is represented as mean percentage of body weight change ± SD. Body weight data was normalized to the initial weight of each animal (*n*=10mice/group) and examined in 2 independent experiments. **g,h.** Data is represented as mean percentage of iRBCs ± SEM, determined in 2-3 independent experiments (*n*=10-15mice/group). **i.** Data is represented as box-whisker plot of mean merozoite number per schizont ± SD (2-way ANOVA), with the median represented at the center line. Boxplots show the data of 3 independent experiments and for the following number of schizonts: SAMS-DD +TMP, *n*=152; SAMS-DD -TMP, *n*=131. **Figure 1 – Figure supplement 2 - Source data 1** Mean fluorescence intensity levels of *P. falciparum* infected RBCs and schizonts developing in MetD conditions related to Figure 1 – Figure supplement 2b,c. **Figure 1 – Figure supplement 2 - Source data 2** Comparison of replication rates in *P. berghei* parasites growing under AA deficient conditions or SAMS knock-down parasites by fluorescence or brightfield microscopy, related to Figure 1 – Figure supplement 2d,e. **Figure 1 – Figure supplement 2 - Source data 3** Mice body weight change upon providing a 2-week MetD regimen, related to Figure 1 – Figure supplement 2f. **Figure 1 – Figure supplement 2 - Source data 4** Parasitemia levels of *P. berghei* parasites growing in MetD media or parasites knock-down for the SAMS enzyme, related to Figure 1 – Figure supplement 2g,h. **Figure 1 – Figure supplement 2 - Source data 5** *In vivo* replication rates of *P. berghei* parasites knock-down for the SAMS enzyme, related to Figure 1 – Figure supplement 2i.

**Figure 2 – Figure supplement 1.**
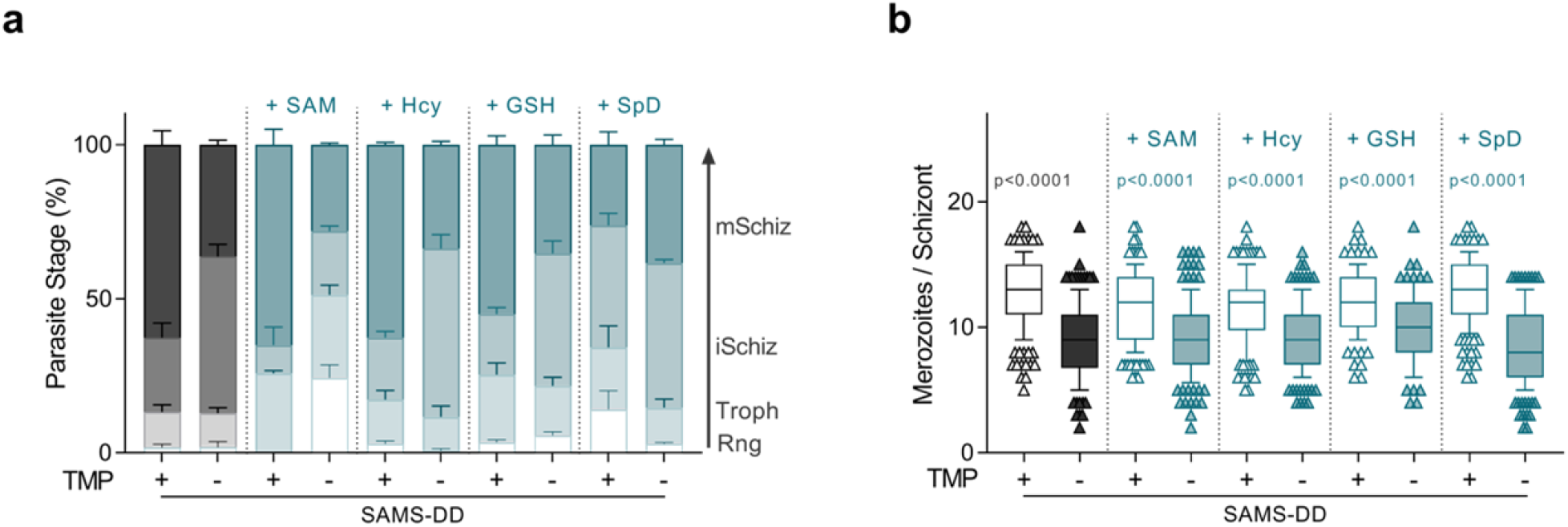
Phenotypic screen of *P. berghei* parasites lacking the SAMS enzyme, upon supplementation with the SAM-downstream metabolites. **(a, b)** Quantification of *P. berghei* **(a)** intra-erythrocytic developmental stages or **(b)** boxplot of mean merozoite numbers in *Pb*SAMS-DD parasites cultured in media supplemented with the SAM-dependent metabolites: 1.5mM S-adenosylmethionine (SAM), 20μM homocysteine (Hcy), 1mM glutathione (GSH) and 1mM spermidine (spD) using the *in vitro* maturation assay. Prior to *in vitro* culture, *Pb*SAMS-DD-infected mice were treated (+TMP), or not (-TMP) with TMP in drinking water. **a.** Parasite staging was performed in triplicate, averaged and repeated in 2-3 independent experiments. Data is shown as mean percentage of parasites at each developmental stage, with error bars representing SEM. **b**. Data is represented as box-whisker plot of mean merozoite number per schizont ± SD (2-way ANOVA), with the median represented at the center line. Boxplots show the data of 3 independent experiments and for the following number of schizonts: SAMS-DD: +TMP, *n*=200; -TMP, *n*=210; +TMP +SAM, *n*=141; -TMP +SAM, *n*=135; +TMP +Hcy, *n*=130; -TMP +Hcy, *n*=129; +TMP +GSH*, n*=102; -TMP +GSH *n*=83; +TMP +SpD, *n*=154; -TMP +SpD *n*=154. **Figure 2 – Figure supplement 1 - Source data 1** Parasite staging analysis of *P. berghei* parasites lacking the SAMS enzyme *in vitro* cultured in media supplemented with the SAM-downstream metabolites, related to Figure 2, Figure supplement 1a. **Figure 2 – Figure supplement 1 - Source data 2** Quantification of parasite replication rates in *P. berghei* parasites lacking the SAMS enzyme, *in vitro* cultured in media supplemented with the SAM-downstream metabolites, related to Figure 2, Figure supplement 1b.

**Figure 3 – Figure supplement 1.**
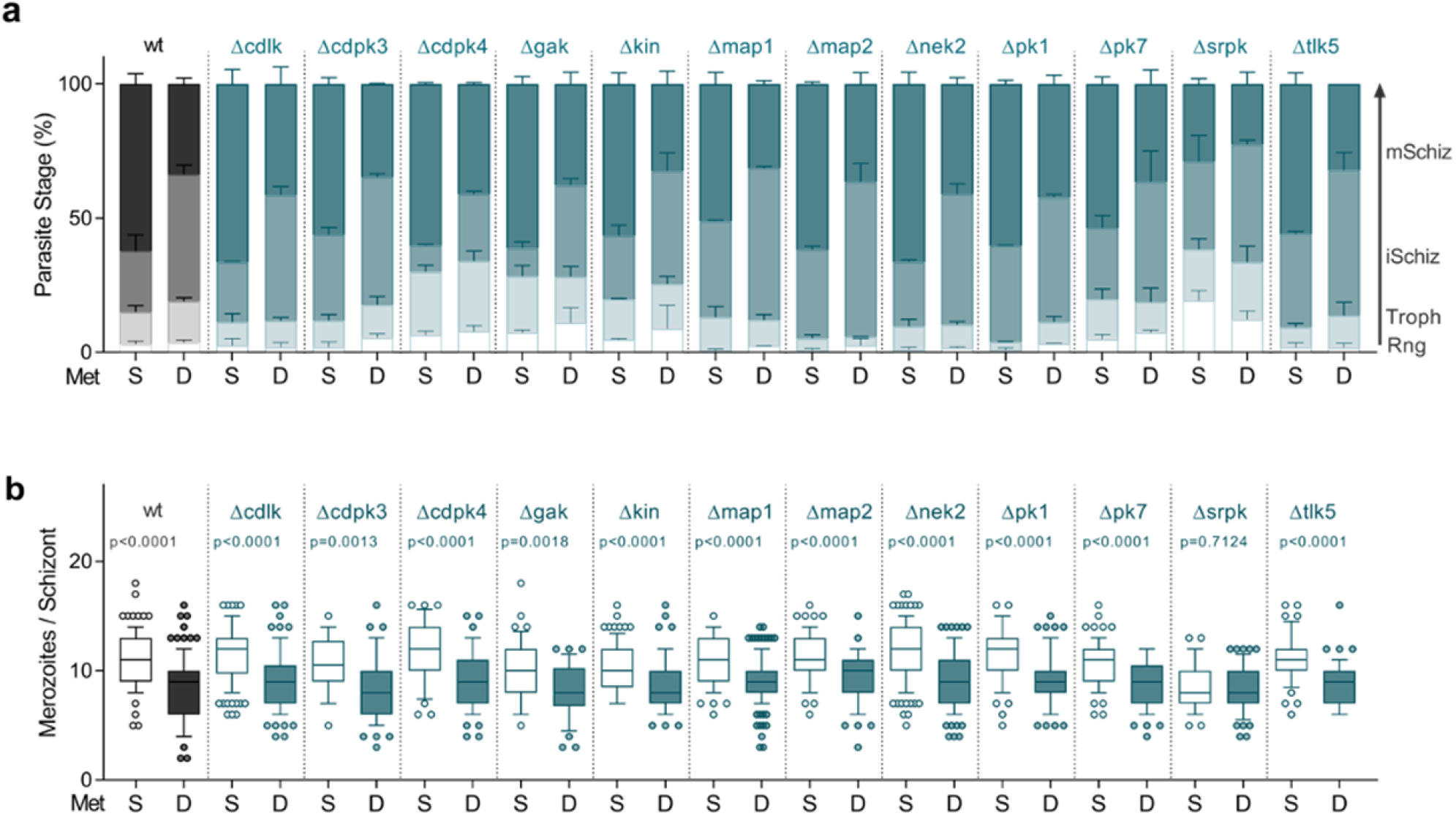
Phenotypic screen of *P. berghei* kinase knockout mutant parasites *in vitro* cultured in MetD media. **(a, b)** Quantification of **(a)** intra-erythrocytic developmental stage progression and **(b)** boxplot of mean merozoite numbers in *P. berghei* wild-type and kinase mutant parasite lines using the *in vitro* maturation assay. **a.** Parasite staging was performed in triplicate, averaged and repeated in 2-3 independent experiments. Data is shown as mean percentage of parasites at each developmental stage, with error bars representing SEM. **b**. Data is represented as box-whisker plot of mean merozoite number per schizont ± SD (2-way ANOVA), with the median represented at the center line. Boxplots show the data of 3 independent experiments and for the following number of schizonts: *P.berghei* wt: MetS, *n*=92; MetD, *n*= 99; Δcdlk: MetS, *n*=110; MetD, *n*=81; Δcdpk3: MetS, *n*= 28; MetD, *n*= 55; Δcdpk4: MetS, *n*=33; MetD, *n*=56; Δgak: MetS, *n*=43; MetD, *n*=34; Δkin: MetS, *n*=85; MetD, *n*=80; Δmap1: MetS, *n*=82; MetD, *n*=133; Δmap2: MetS, *n*=64; MetD, *n*=58; Δnek2: MetS, *n*=112; MetD, *n*=96; Δpk1: MetS, *n*=49; MetD, *n*=64; Δpk7: MetS, *n*=69; MetD, *n*=73; Δsrpk: MetS, *n*=37; MetD, *n*=54; Δtlk5: MetS, *n*=44; MetD, *n*=51. **Figure 3 – Figure supplement 1 - Source data 1** Parasite staging analysis of *P. berghei* kinase knockout mutants *in vitro* cultured in MetD media, related to Figure 3, Figure supplement 1a. **Figure 3 – Figure supplement 1 - Source data 2** Quantification of parasite replication rates in *P. berghei* kinase knockout mutants, *in vitro* cultured in MetD media, related to Figure 3, Figure supplement 1b.

**Figure 4 – Figure supplement 1.**
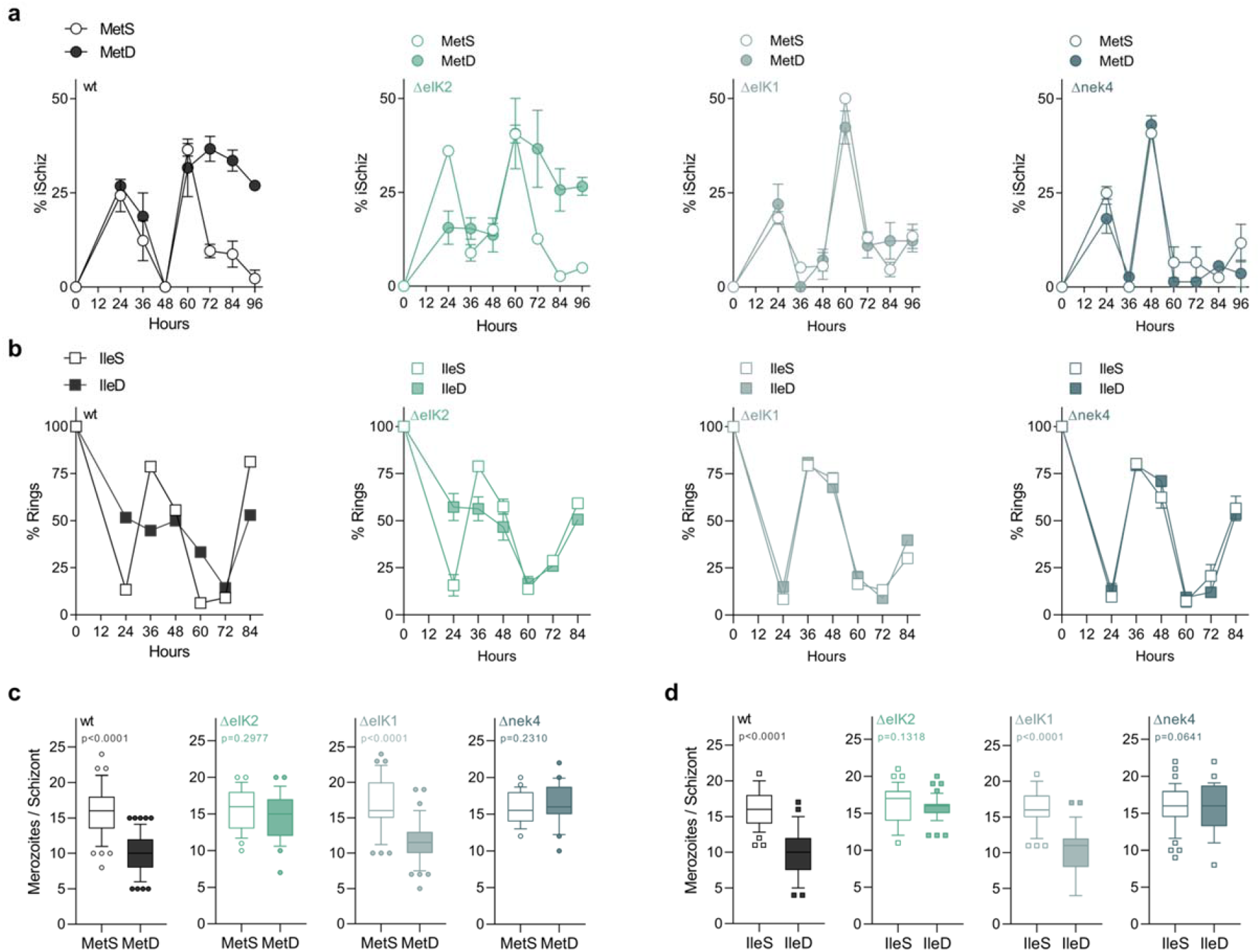
AA depletion impact in the human malaria spp. *P. falciparum*. **(a)** Time course analysis of immature schizont-stage *P. falciparum* 3D7 WT and kinase mutant parasites cultured in MetS- or MetD-media (wt, black; ΔeIK2, emerald green; ΔeIK1, light teal; Δnek4, dark teal). **(b)** Time course analysis of ring-stage *P. falciparum* 3D7 WT and kinase mutant parasites cultured in IleS- or IleD-media. **(c, d)** Boxplot of mean merozoite numbers in *P. falciparum* 3D7 WT and kinase mutant parasites, cultured in **(c)** MetD or **(d)** IleD conditions. **a,b.** Data represents the mean percentage of **(a)** immature schizonts or **(b)** ring-stages ± SEM, each performed at least in triplicate, averaged and determined in 2 independent experiments. **c,d.** Data is represented as box-whisker plot of mean merozoite number per schizont ± SD (Mann-Whitney), with the median represented at the center line. Boxplots show the data of 2 independent experiments and for the following number of schizonts: wild-type: MetS, *n*= 57; MetD, *n*= 58, ΔeIK2: MetS, *n*= 26; MetD, *n*= 25; ΔeIK1: MetS, *n*= 35; MetD, *n*= 44; and Δnek4: MetS, *n*= 22; MetD, *n*= 20; (d) IleS, *n*= 37; IleD, *n*= 33; ΔeIK2: IleS, *n*= 37; IleD, *n*= 42; ΔeIK1: IleS, *n*= 41; IleD, *n*= 36; and Δnek4: IleS, *n*= 45; IleD, *n*= 28. **Figure 4 – Figure supplement 1 - Source data 1** Time course analysis of the percentage of immature schizonts produced by *P. falciparum* wt and the kinase mutant knockout lines, ΔeIK1, ΔeIK2 and Δnek4 under Met deficient conditions, related to Figure 4, Figure supplement 1a. **Figure 4 – Figure supplement 1 - Source data 2** Time course analysis of the percentage of *P. falciparum* ring-stage parasites in wt and the kinase mutant knockout lines, ΔeIK1, ΔeIK2 and Δnek4 under Ile deficient conditions, related to Figure 4, Figure supplement 1b. **Figure 4 – Figure supplement 1 - Source data 3** Replication rates of *P. falciparum* wt and the kinase mutant knockout lines, ΔeIK1, ΔeIK2 and Δnek4 developing under AA deficient conditions, related to Figure 4, Figure supplement 1c,d.

| Item | Recommendation | Section/line number, or reason for not reporting |
| --- | --- | --- |
| **Study design** | 1 For each experiment, provide brief details of study design including:<br>The groups being compared, including control groups. If no control group has been used, the rationale should be stated.<br>The experimental unit (e.g. a single animal, litter, or cage of animals). | a,b.Methods:"Mice, diets and treatments"<br>Lines 396-401 |
| **Sample size** | 2<br>Specify the exact number of experimental units allocated to each group, and the total number in each experiment. Also indicate the total number of animals used.<br>Explain how the sample size was decided. Provide details of any *a priori* sample size calculation, if done. | a,b. Methods: "Statistical Analysis"<br>Lines 584-588 |
| **Inclusion and exclusion criteria** | 3<br>Describe any criteria used for including and excluding animals (or experimental units) during the experiment, and data points during the analysis. Specify if these criteria were established *a priori*. If no criteria were set, state this explicitly.<br>For each experimental group, report any animals, experimental units or data points not included in the analysis and explain why. If there were no exclusions, state so.<br>For each analysis, report the exact value of *n* in each experimental group. | a,b.No data points were excluded from analysis<br>c. Described in respective Figure Legend<br>Methods: Lines 585-588 |
| **Randomisation** | 4<br>State whether randomisation was used to allocate experimental units to control and treatment groups. If done, provide the method used to generate the randomisation sequence.<br>Describe the strategy used to minimise potential confounders such as the order of treatments and measurements, or animal/cage location. If confounders were not controlled, state this explicitly. | a,b.Methods:"Mice, diets and treatments"<br>Lines 396-401; 444-446; 491-495 |
| **Blinding** | 5 Describe who was aware of the group allocation at the different stages of the experiment (during the allocation, the conduct of the experiment, the outcome assessment, and the data analysis). | Investigators were not blinded to the group allocation. Lines 400-402 |
| **Outcome measures** | 6<br>Clearly define all outcome measures assessed (e.g. cell death, molecular markers, or behavioural changes).<br>For hypothesis-testing studies, specify the primary outcome measure, i.e. the outcome measure that was used to determine the sample size. | a,b. Methods section<br>a,b. Methods titles |
| **Statistical methods** | 7<br>Provide details of the statistical methods used for each analysis, including software used.<br>Describe any methods used to assess whether the data met the assumptions of the statistical approach, and what was done if the assumptions were not met. | Methods: "Statistical Analysis"<br>Lines 579-588 |
| **Experimental animals** | 8<br>Provide species-appropriate details of the animals used, including species, strain and substrain, sex, age or developmental stage, and, if relevant, weight.<br>Provide further relevant information on the provenance of animals, health/immune status, genetic modification status, genotype, and any previous procedures. | a. Methods: Lines 395-406<br>b. Methods: Lines 437-446 |
| **Experimental procedures** | 9 For each experimental group, including controls, describe the procedures in enough detail to allow others to replicate them, including:<br>What was done, how it was done and what was used.<br>When and how often.<br>Where (including detail of any acclimatisation periods).<br>Why (provide rationale for procedures). | a-d.Methods:<br>Lines 395-406; 410-435<br>Lines 437-446; 475-480 |
| **Results** | 10 For each experiment conducted, including independent replications, report:<br>Summary/descriptive statistics for each experimental group, with a measure of variability where applicable (e.g. mean and SD, or median and range).<br>If applicable, the effect size with a confidence interval. | Described in respective Figure Legend |

